# Single cell characterization of B-lymphoid differentiation and leukemic cell states during chemotherapy in ETV6-RUNX1 positive pediatric leukemia identifies drug-targetable transcription factor activities

**DOI:** 10.1101/2020.05.27.116293

**Authors:** Juha Mehtonen, Susanna Teppo, Mari Lahnalampi, Aleksi Kokko, Riina Kaukonen, Laura Oksa, Maria Bouvy-Liivrand, Alena Malyukova, Saara Laukkanen, Petri I. Mäkinen, Samuli Rounioja, Pekka Ruusuvuori, Olle Sangfelt, Riikka Lund, Tapio Lönnberg, Olli Lohi, Merja Heinäniemi

## Abstract

Tight regulatory loops orchestrate commitment to B-cell fate within bone marrow. Genetic lesions in this gene regulatory network underlie the emergence of the most common childhood cancer, acute lymphoblastic leukemia (ALL). The initial genetic hits, including the common translocation that fuses ETV6 and RUNX1 genes, lead to arrested cell differentiation. Here, we aimed to characterize transcription factor activities along the B-lineage differentiation trajectory as a reference to characterize the aberrant cell states present in leukemic bone marrow, and to identify those transcription factors that maintain cancer-specific cell states for more precise therapeutic intervention.

We compared normal B-lineage differentiation and *in vivo* leukemic cell states using single cell RNA-sequencing (scRNA-seq) and several complementary genomics profiles. Based on statistical tools for scRNA-seq, we benchmarked a workflow to resolve transcription factor activities and gene expression distribution changes in healthy bone marrow lymphoid cell states. We compared these to ALL bone marrow at diagnosis and *in vivo* during chemotherapy, focusing on leukemias carrying the ETV6-RUNX1 fusion.

We show that lymphoid cell transcription factor activities uncovered from bone marrow scRNA-seq have high correspondence with independent ATAC- and ChIP-seq data. Using this comprehensive reference for regulatory factors coordinating B-lineage differentiation, our analysis of ETV6-RUNX1-positive ALL cases revealed elevated activity of multiple ETS-transcription factors in leukemic cells states, including the leukemia genome-wide association study hit ELK3. The accompanying gene expression changes associated with natural killer cell inactivation and depletion in the leukemic immune microenvironment. Moreover, our results suggest that the abundance of G1 cell cycle state at diagnosis and lack of differentiation-associated regulatory network changes during induction chemotherapy represent features of chemoresistance. To target the leukemic regulatory program and thereby overcome treatment-resistance, we show that selective inhibitors of ETS-transcription factors could effectively reduce cell viability.

Our data provide a detailed picture of the transcription factor activities that characterize both normal B-lineage differentiation and those acquired in leukemic bone marrow and provide a rational basis for new treatment strategies targeting the immune microenvironment and the active regulatory network in leukemia.

## Background

Failures in lymphoid cell differentiation underlie the emergence of acute lymphoblastic leukemia (ALL) that peaks in incidence in childhood [1]. Recently, 35 potential cell states in hematopoiesis were resolved using single-cell RNA-seq (scRNA-seq) data based on eight healthy bone marrow (BM) donors profiled by the Human Cell Atlas (HCA) groups, comprising approximately 100 000 cells [2]. Understanding normal B cell differentiation in BM forms the basis to characterize the aberrant cell states in cancers that originate from lymphoid progenitor cells. Previous work has identified tight regulatory loops that orchestrate B-cell fate [3]. However, their activity along the single cell resolution trajectory in human B-lineage has not been studied in detail.

The genetic basis of ALL initiation and progression is mechanistically linked to alterations in key lymphoid transcription factors (TFs) [1]. The most common translocation t(12;21) generates a fusion between two transcription factors (TFs): the repressive domain of ETV6 is fused with RUNX1, retaining the RUNT-DNA-binding domain. This confers cells with functional properties that sustain self-renewal and survival [4]. We and others have shown that the aberrant ETV6-RUNX1 (E/R) TF-fusion can silence key genes and regulatory regions [5–9]. In effect, cells become arrested at a lymphoid progenitor state [7,10], whereby additional DNA lesions can accumulate, which especially in E/R-leukemias are driven by a transcription-coupled mechanism that results in off-targeting of the recombination activating genes (RAG) complex [11,12]. However, the emerging cell state heterogeneity that manifests at diagnosis and during chemotherapy within the bone marrow remains poorly characterized.

In the clinics, the accumulated knowledge regarding initiating genetic lesions has been implemented into diagnostic screens that inform choices between chemotherapy regimes that differ in intensity. However, almost half of relapses occur in children presenting initially with good-risk cytogenetic features such as E/R [13], thus raising the question what underlies their resistance. Epigenetic changes driven by TF, coregulator and chromatin modifier activities in the blast cells contribute to the blast cell phenotype [14,15]. The epigenetic plasticity of leukemic cells may support resistant states [16,17], allow conversion into quiescent stem-like states or lineage switching to escape the cytotoxic agents [18–22]. This poses a challenge in the design of drug therapy and urges the development of new therapies informed by characterization of the cancer cells and their cross-talk with the microenvironment.

Single cell genomics holds promise to resolve the leukemic gene regulatory programs even in small cell populations, based on mRNA, chromatin and DNA profiles [23]. Computational analysis can resolve TF activity, transcriptome dynamics and capture changes in gene expression distributions between cell states analyzed [24–26]. Here, we set out to elucidate cell states and TF activities characteristic of normal B-lineage differentiation from hematopoietic stem cells (HSC) and to compare these to the E/R+ ALL cases at diagnosis and during standard chemotherapy.

## Results

### Bone marrow B-lineage differentiation states are captured in single cell transcriptomes

For a refined view on early B-cell differentiation, we processed BM scRNA-seq data available from HCA [27] and projected each cell into a two-dimensional map using UMAP (Fig. S1). A branching map centered at *CD34+* HSC was obtained, where cycling progenitor cell states lead to more differentiated cells that predominantly existed in the G1 cell cycle state based on cell cycle marker gene scoring (Fig. S1a-b), while stromal cells or mature T (*CD3D*+), NK (*GNLY*+) and plasma B cells, which mature outside the bone marrow, clustered separately (Fig. S1c).

We separated the B-lineage branch for further analysis, resulting in a reference dataset for B-lineage differentiation from HSCs with 11 clusters (Fig. 1a). The first two clusters corresponded to HSC (in G1 or cycling cell cycle states S/G2/M). *DNA nucleotidylexotransferase (DNTT*, also known as TdT) and *MME* (also known as CD10) marker gene expression distinguishes the early lymphoid progenitors (CLP, cluster 3) that progress into the CD19-expressing cycling and G1 pro-B cell states (Fig. 1b). Furthermore, three pre-B cell clusters (lacking *DNTT* expression) segregated on the map, corresponding to the cycling large pre-B state, followed by pre-B-I and pre-B-II cells in the G1 cell cycle state. The pre-B-II and the subsequent immature B cell clusters were defined by MS4A1(CD20)-positivity [28,29]. The pseudo-temporal ordering of the clusters, based on diffusion pseudotime analysis, is shown in Figure 1c. The progression between cell states based on this analysis is in agreement with the assigned differentiation stages. These cell state annotations had high agreement also with differentiation state scoring using gene sets defined by flow-sorted B-cell populations (Fig. 1d) [30]. However, these gene sets defined from bulk transcriptomes scored highly only in the cycling cell states. Therefore, we additionally distinguished marker genes for each cluster from the single cell analysis (Table S1) to facilitate BM B-lineage cell state assignment in future studies.

**Figure 1.**
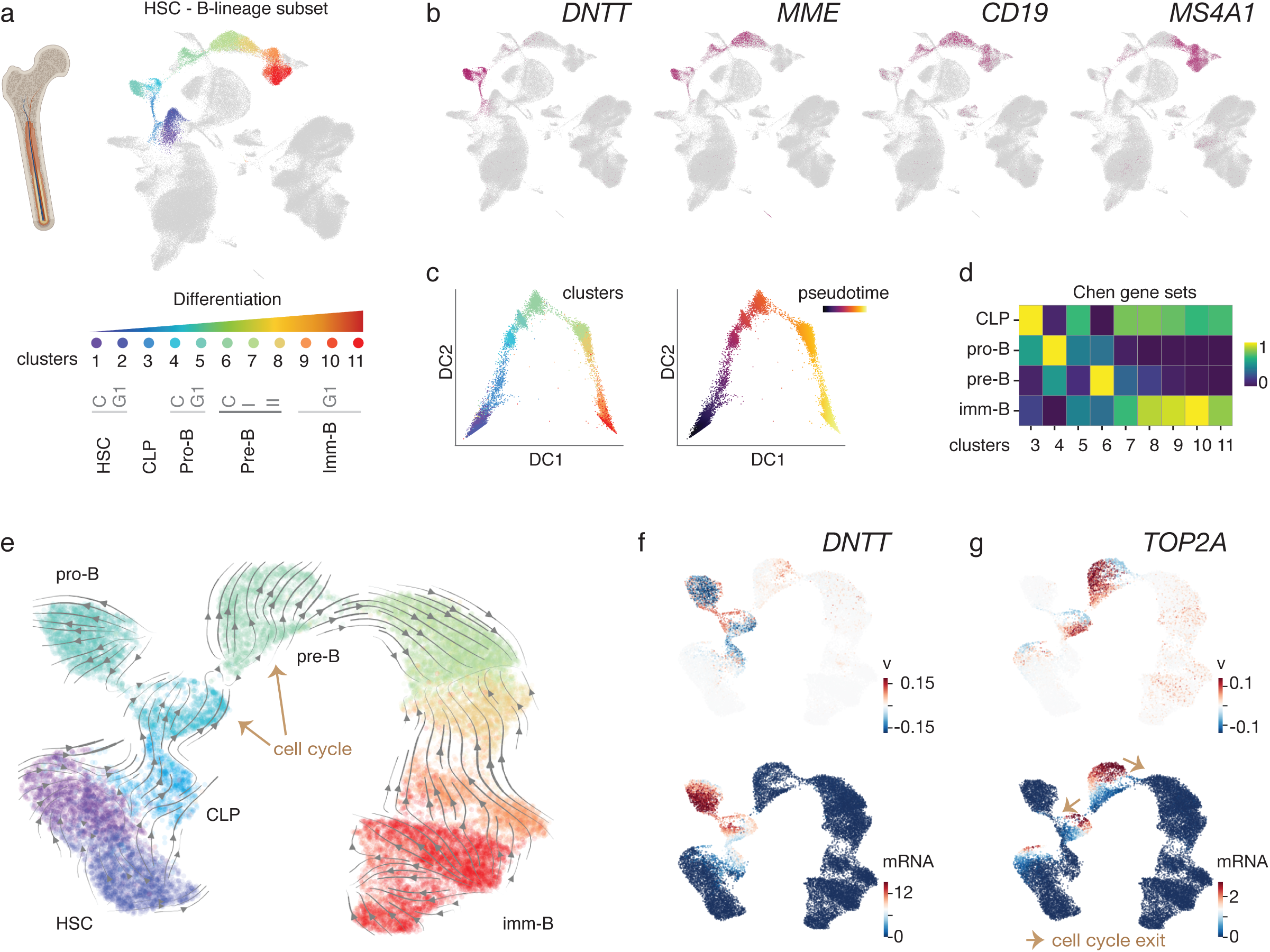
B-lymphoid differentiation states separate in bone marrow scRNA-seq. a. scRNA-seq clusters for the B-lymphoid lineage defined from HCA BM scRNA-seq data are shown in color on the UMAP visualization and annotated by differentiation stage (refer to Fig. S1 for other cell type annotations). b. Marker gene expression level is colored on the UMAP, where darker tones of red indicate high expression. c. Diffusion pseudotime ordering of cells is shown with colors corresponding to clusters shown in C (left) or pseudotime (right). d. Scores for gene sets corresponding to distinct B-cell differentiation states[30] are visualized as a heatmap. e. Differentiation dynamics based on RNA velocyto analysis is shown for the B-lymphoid cell states. Arrows correspond to predicted direction of cell state transitions. f and g. RNA velocities (top panel) compared to spliced mRNA counts (bottom panel) of the early B-cell marker gene *DNTT* and the G2/M-phase specific gene *TOP2A* are shown. Red tones correspond to high velocity or mRNA level, respectively.

To delineate the gene expression changes that characterize the cell state transitions in early B-lineage differentiation, we compared the cell clusters sequentially along the pseudotime trajectory (HSC -> CLP -> pro-B -> pre-B -> immature B cell state). Using the scDD tool [26], changes in mRNA detection (as proportion of zeros), differences in mean expression and modality could be distinguished for 2201 genes in total (Fig. S2; Table S2). Analysis of the RNA dynamics of this gene cohort based on RNA velocity [25,31] allowed further resolving the B-lineage cell state map (Fig. 1e). In this analysis, both spliced and unspliced counts are used to estimate the velocity of gene expression change, thus extending the cell state representation with gene regulatory dynamics (see Methods). This is illustrated by *DNTT* (Fig. 1f) that is first upregulated (red tones correspond to positive velocity, top panel) in early lymphoid cells and further increases in mRNA expression (red tones indicating high spliced mRNA counts, bottom panel) at the pro-B state. The pro-B G1 cell state separates as a branch in these analyses, indicating the possibility that this cell state is present as a progenitor pool. Moreover, two successive cycling cell states precede the cell cycle exit into the small pre-B state: the S-phase marker *PCNA* is upregulated (positive velocity) as cells progress from early lymphoid to the first cycling state (pro-B cycling) (Fig. S1d) and its mRNA peaks at S-phase cells, coinciding with increasing *TOP2A* velocity (Fig. 1g, top panel, G2/M marker gene) that subsequently peaks in its mRNA level at the G2/M state. The successive increases in the velocity and mRNA levels of these cell cycle state markers indicate the direction of cells on the map and the final exit from the cell cycle into pre-B I G1 state (Fig. 1g, lower panel).

### TF activity changes reveal the regulatory dynamics of B cell differentiation

The cell state transitions along the B-lineage trajectory are tightly controlled by TFs. To characterize TF, coregulator (CR), chromatin modifier (CM) and splicing/transcription complex (ST) activities at fine-resolution, we performed discovery of so-called TF regulons with a workflow based on the SCENIC tool [24] (see Methods for details). Significant predictors for cell states were analyzed by linear model fitting using regulons that were reproducibly identified across training and test set splits. The regulon activity scoring across the B-lineage differentiation stages is shown in Figure 2a (Table S3) for regulons passing a stringent R^2^ cut-off (0.5). Expression levels for TFs involved in the main B-lineage commitment loop (B-lineage TFs reviewed in [32,33]) are shown for comparison in Figure 2b. EBF1, FOXO1, LEF1 and TCF4, together with ETS-factors ERG and FLI1, displayed the highest activity (in red) in pro-B cells in our analysis, while TCF3 and PAX5 had similarly high activity in both pro- and pre-B cell states.

SPIB and IRF4 activity was elevated later at pre-B cells, together with several negative regulons for TFs with known repressive function such as BCL11A and known co-repressor complex components HDAC2 and TBL1XR1 that interact with glucocorticoid receptor to promote terminal differentiation.

As independent validation, we first retrieved bulk ATAC-seq profiles from pro-B cells [34]. Significantly enriched TF motifs confirmed 9/12 TF regulons (EBF1, FOXO1, TCF3, RFX5, IRF1, TCF4, LEF1, ERG, FLI1) that our analysis associated with the pro-B G1 cell state (Fig. 2c). Next, we examined closer the regulon gene sets that include TF targets discovered based on TF-to-target gene expression correlation and TF-motif analysis at each target gene locus. We categorized the predicted targets based on how many training/test set splits supported them in the regulon discovery phase. To test whether the predicted targets were bound by the TF, we retrieved ChIP-seq data for PAX5, EBF1 and BCL11A, available in the human cell line model Nalm-6 (see Methods). Peak to gene associations were obtained using the tool GREAT [35], and compared to SCENIC predictions (Table S3). For PAX5 and EBF1, over 75% of predicted targets had a ChIP-seq peak association (Fig. 2d). The validation for the BCL11A repressive regulon was initially low (< 25%). However, upon modification of the regulon discovery strategy (see Methods; data shown in Fig. 2a corresponds to updated regulon discovery), we could improve this nearly 2-fold. Moreover, targets discovered across multiple training data splits (Npred, number of iterations supporting the target) were associated with more ChIP-seq peaks (Fig. 2e, Fig. S3), including the most prominent peaks based on ChIP peak score (Fig. 2f, low ranks corresponds to best ChIP scores). The number of associated peaks and their relative peak ranking is further illustrated for top 50 genes from the PAX5 regulon (Fig. 2g, targets ranked by Npred). ChIP-seq validated genes include known PAX5 targets from confirmed regulatory loops *(EBF1, IRF4, BACH2)* and B-cell maturation pathways [36–38]. The high agreement of ATAC-seq motif enrichment and the verified TF binding at target gene set loci based on ChIP-seq provides evidence that the TF activity scoring reflects bona fide active regulatory interactions.

**Figure 2.**
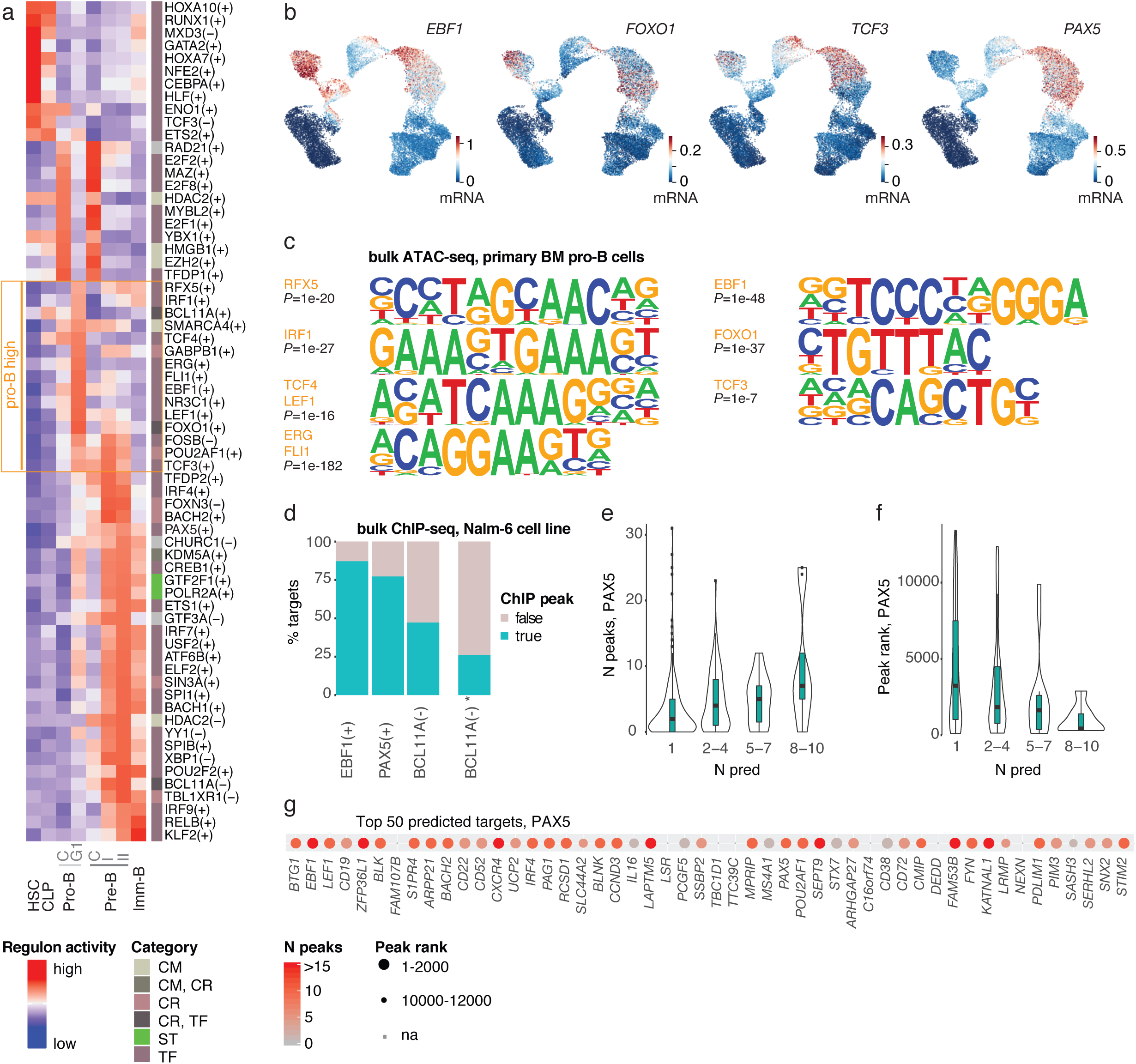
Transcription factor activities across B-lineage differentiation. a. Regulon activity score is visualized as a heatmap (tones of red indicate high activity). The annotated functional category (CM=chromatin modifier; CR=coregulator; TF=transcription factor; ST=splicing/transcription complex) of each regulon is shown. b. Gene expression levels for the TFs *EBF1, FOXO1, TCF3* and *PAX5* are indicated in color on the B-lineage scRNA-seq map. c. Significant motifs matching pro-B active TFs (indicated in panel a) are shown from pro-B cell bulk ATAC-seq. d. Percentage of TF regulon target genes associated with ChIP-seq peaks is shown for EBF1(+), PAX5(+) and BCL11A(-) regulons obtained with the customized workflow. BCL11A(-)* corresponds to the initial regulon discovered by default SCENIC run. e. The distribution of ChIP-seq peaks associated to targets is shown as a combined violin and box plot for predicted targets from PAX5(+) regulon. Targets are binned based on the number of training set runs supporting each target (Npred). The y-axis corresponds to 0.05-0.95 quantile range. f. Similarly as in e, the ChIP-seq peak rank distribution is visualized across binned PAX5(+) regulon genes. Lower rank corresponds to higher ChIP peak score. g. The ChIP peak data is visualized using a dot plot for top 50 predicted PAX5 targets. The color corresponds to the number of associated peaks and the dot size indicates binned ChIP-peak rank (bin size 2000).

In summary, our analysis of healthy BM single cell transcriptomes provides a comprehensive reference for gene expression and TF activity changes that characterize early B-lineage differentiation at single cell resolution.

### E/R-leukemic cells resemble the pro-B cell state and display heterogeneity in cell cycle activity

Lymphoblastic leukemias arise as a consequence of arrested cell differentiation and often carry initiating genetic lesions directly affecting key lymphoid TFs. To characterize leukemic cells carrying the most common TF fusion in ALL (E/R), we performed scRNA-seq on six pediatric E/R+ pre-B-ALL cases, collecting from each the diagnostic BM and from two cases BM at day 15 during induction chemotherapy (Fig. 3a, Table 1). The leukemic cell clusters in each donor were identified based on *DNTT* expression and their clear separation from normal bone marrow cell types (Fig. 3b) (Fig. S4a-c). The cycling leukemic states clustered across donors directly, while the similarity of G1 leukemic cells could be ascertained by correcting for donor effect (Fig. S4d). Based on the B-lineage cluster-specific gene sets, the diagnostic leukemic blasts resembled the pro-B differentiation state (Fig. 3c). This analysis was supported by label transfer analysis using Seurat [39] (Fig. S4c) that similarly identified pro-B cells as the closest normal differentiation state, in agreement with previous studies [4,7,40]. The cell cycle state distribution differed between cases, from lowest proportion of cycling cells in ALL3 to highest in ALL9 (Fig. 3d). For the two cases (ALL10 and ALL12) with mid-induction therapy BM profiles, the cells collected at day 15 separated as distinct cell states (Fig. 3a and d), indicating that treatment further alters leukemic cell states.

**Figure 3.**
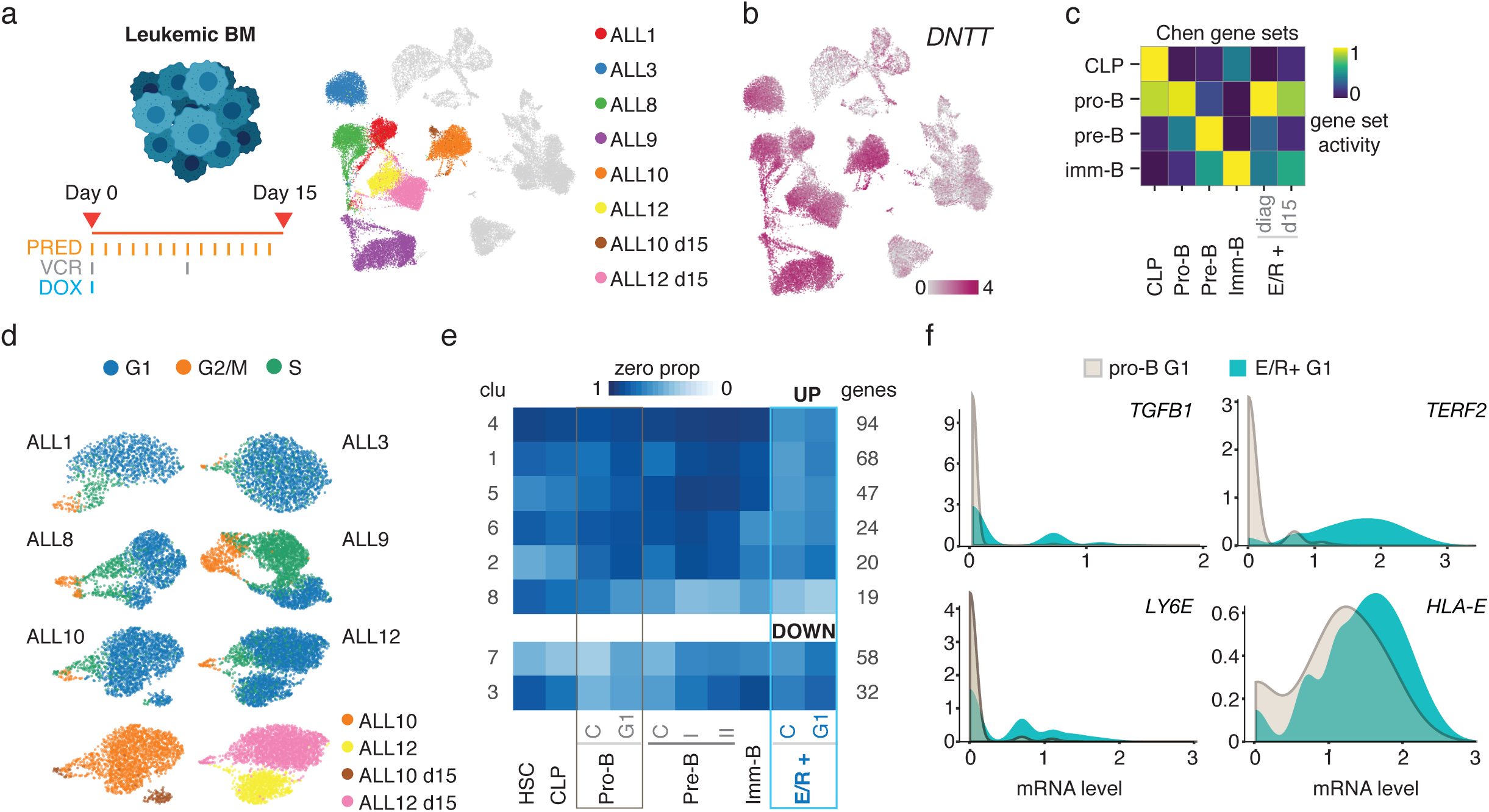
Comparison of E/R+ cells to normal pro-B cells. a. Six diagnostic and two post-treatment bone marrow samples analyzed with scRNA-seq are shown on the UMAP representation. b. Expression level of the *DNTT* marker gene is shown in color on the ALL BM UMAP. c. Gene set scoring of differentiation stage is shown as a heatmap, comparing ALL cells to normal bone marrow lymphoid cells. d. Computationally predicted cell cycle state of leukemic cells is colored separately for each donor on a UMAP. For ALL10 and ALL12 the sample origin (diagnostic or day 15 post-treatment) is indicated in the bottom panel. e. Eight clusters formed from genes that distinguish ALL cells from normal pro-B cells are shown in the heatmap. The data corresponds to cluster centroids and the colors indicate the mRNA detection metric ZP, with dark blue tones indicating low expression (high ZP) and light tones corresponding to larger proportion of cells expressing the genes in each cluster. The number of genes in each cluster is indicated on the right. f. Genes modulating NK cell activity *(TGFB1, TERF2, LY6E* and *HLA-E)* plotted as density plots that compare the gene expression distribution of E/R+ cells to pro-B cells (both in G1 cell cycle state).

**Table 1.** Leukemic blast percentage during induction therapy determined by BM flow cytometry. E/R-positive patient samples used in genomics experiments are shown.

| **Sample ID** | **Leukemic blast-% at diagnosis** | **Leukemic blast-% at day 15** | **Leukemic blast-% at day 29** |
| --- | --- | --- | --- |
| ALL1 | 94 | 10 | 0.3 |
| ALL3 | 95 | 74 | 0.16 |
| ALL8 | 93 | 0.93 | 0.02 |
| ALL9 | 79 | 0.17 | 0.01 |
| ALL10 | 65 | 10 | 0.08 |
| ALL12 | 90 | 59 | 0.2 |
| ALL7 | 27 | 0 | 0 |
| ALL13 | 80 | 0.04 | 0 |

Next, we aimed to further characterize how the diagnostic E/R leukemic cells differ from pro-B cells by comparing separately the gene expression distributions of cycling and G1 state cells to normal pro-B cells. For the majority of genes, the most notable change upon normal B-lineage differentiation was in the zero proportion (ZP) metric that captures the fraction of cells with zero counts for a gene of interest, as exemplified for top 50 genes up- and downregulated in pro-B to pre-B transition (Fig. S2). Therefore, we used ZP for clustering the 272 up- and 90 downregulated genes found in both G1 and cycling cell state comparisons of E/R+ and pro-B cells (Fig. 3e; Table S2). Compared to other cell states along the B-lineage differentiation trajectory, about one-third of the upregulated genes were expressed at the highest level in E/R+ cells (cluster 4), while genes in clusters 1, 2, 5 and 6 showed expression in leukemia and normal stem/progenitor cells (Fig. 3e). A smaller fraction (19 genes, cluster 8) were highly expressed in normal pre- or immature B-cells. Considering that some gene expression patterns resembled the pre-B cell state, yet the leukemic cells appeared arrested at the pro-B state, we further identified genes that are normally regulated in the pro-B to pre-B transition, to distinguish additional genes associated with the differentiation arrest. In total 97 genes normally upregulated upon transition to pre-B state remained at similar low expression level as in normal pro-B cells, while 145 genes downregulated during differentiation remained expressed in leukemic (Table S2).

Pathway enrichment analysis (Table S4) revealed that several of the upregulated genes associated with cytokine, chemokine and growth factor pathways, in particular those involved in the negative regulation of NK cell-mediated cytotoxicity. Previous study in ALL implicated elevated TGF-β expression in immune evasion [41]. *TGFB1* and three additional genes, *LY6E, TERF2* and *HLA-E*, contributing to lower NK cell recruitment and activation [42—44] were found up-regulated in our analysis comparing the expression distribution of E/R+ G1 cells to pro-B G1 cells (Fig. 3f).

### The E/R+ BM immune microenvironment has low abundance and activity of NK cells

The increase in cells expressing genes that may suppress NK cell activity prompted further analysis of the BM immune cells. In accordance, *GNLY* or *NKG7* positive NK cell numbers were markedly reduced in E/R+ BM compared to HCA BM donors (Fig. 4a). To characterize the immune cell populations further, we combined T and NK cells across HCA and E/R+ ALL donors for joint analysis.

**Figure 4.**
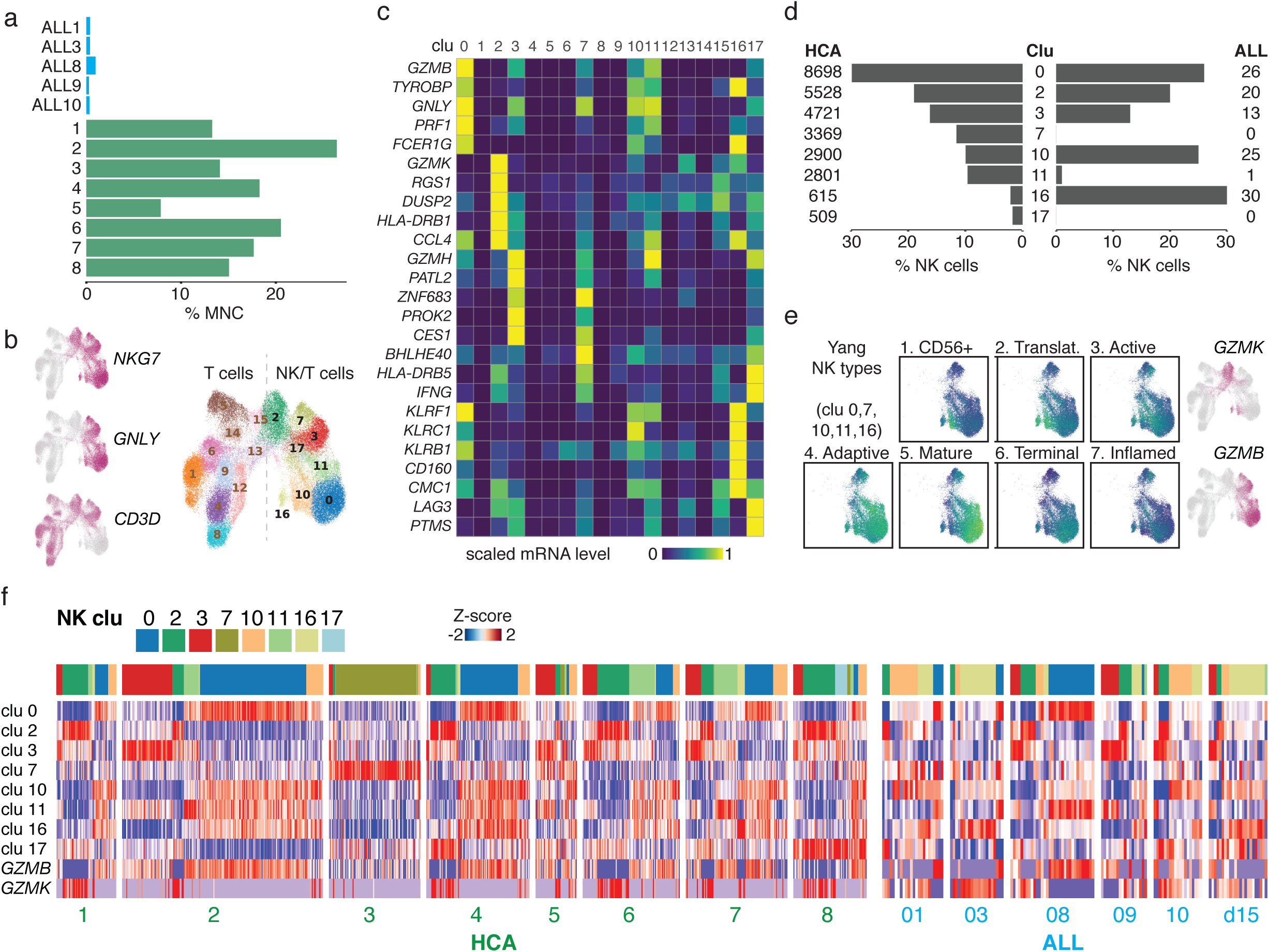
NK cell numbers and activity are low in E/R+ BM. a. Percentage of NK cells in BM scRNA-seq data is shown as barplots across the five diagnostic E/R+ (top panel, in blue, ALL12 samples enriched for B-cells are not shown) and eight normal BM samples (bottom panel, in green). b. Gene expression level for NK and T cell markers and Louvain clustering of cells is shown on the HCA and ALL NK/T cell UMAP. c. Marker genes for NK clusters analyzed are shown as a heatmap. Bright yellow color tones correspond to high expression. d. The percentage (in barplot) and NK cell counts in each cluster are shown from HCA (left) and ALL (right) samples. e. Gene set scores corresponding to scRNA-seq based subtypes[45] are colored on the NK/T cell UMAP. Only cell clusters with low/negative *CD3D* expression used in this comparison are shown. f. NK cell data plotted separately by donor, indicating the cluster assignment for each cell (colored bar above). The heatmap shows the scaled gene set score for NK cell clusters and gene expression for mature vs immature cell markers GZMB and GZMK, respectively. The cells (in columns) are clustered based on gene set activity.

Based on clustering and marker gene analysis, several different NK cell types could be distinguished (Fig. 4b and c). We focused on clusters expressing *GNLY* or *NKG7* (clusters 0, 2, 3, 7, 10, 11, 16) and noticed that the NK cells from ALL BM were disproportionately assigned to these clusters compared to NK cells from HCA donors (Fig. 4d). Specifically, ALL NK cells mainly represented clusters 10 and 16 that matched granzyme K *(GZMK)* expressing immature CD56 bright and translational NK cells (gene set scores in Fig. 4e represent the NK subtypes from a scRNA-seq study[45]). In comparison, the majority of the normal BM NK cells represented the mature or terminal NK cells (cluster 0) that express granzyme B *(GZMB)* and perforin *(PRF1).* Therefore, E/R+ leukemic cells may actively evade NK cell cytotoxicity. However, the frequency of NK types varied between donors (Fig. 4f). Cluster 7 that expressed *IFNG* at high level corresponded almost exclusively to HCA donor 3 and the highly cell cycle active ALL8 and ALL9 resembled more the mature or active NK profile in normal BM compared to other ALL cases.

Taken together, the leukemic cell states differed from normal pro-B differentiation state based on high expression of stem/progenitor cell-specific genes and several immunomodulatory genes. The changes in immunomodulatory genes were reflected as more immature NK cell types within the E/R+ BM.

### The leukemic regulatory program reveals cell state infidelity in TF activities and includes leukemia risk genes

To further decipher the aberrant TF activities contributing to the epigenetic reprogramming that distinguishes E/R+ leukemic cells from normal lymphoid cell states, we repeated the TF regulon activity analysis including the diagnostic leukemic cell states from patient BM (Fig. 5a) (Table S3). Two-third of the regulons passing the linear model fit (R^2^>0.5) were active in pro-B cells and showed elevated activity in E/R+ cells, including several ETS-factors (ELK3, ERG, FLI1), FOXO1, MAX, MAZ, SP4, TCF4 and THAP11. However, our analysis also revealed high activity of RFX5 and NFYC in E/R+ blasts that typically would peak only at the immature B-cell state. This infidelity in differentiation-stage timed TF activities is also manifest in the misexpression of GATA2 that is normally confined to HSC and erythroid progenitors. Furthermore, high but more variable levels of IRF-, KLF-STAT- and CREB1 activity characterized E/R+ cells. Regulons showing diminished activity included RUNX1, SPIB, TCF3 and IRF4 (Fig. 5a).

**Figure 5.**
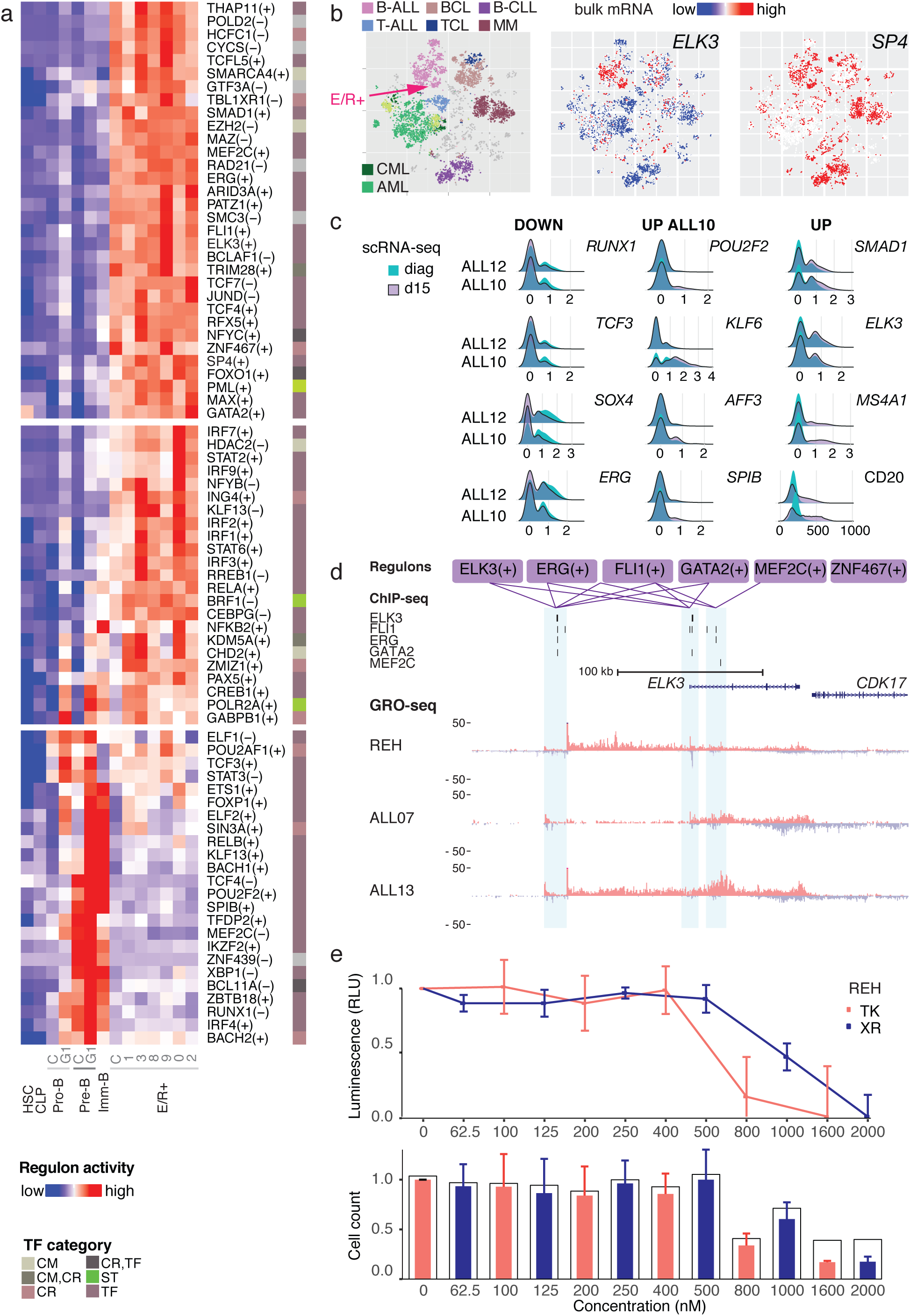
TF activity in E/R+ leukemic cells. a. Regulon activity is visualized as a heatmap as in Fig. 2a comparing E/R+ cells and normal BM cells. b. Bulk mRNA expression data for *ELK3* and *SP4* from Hemap is shown on a t-SNE map comparing transcriptomes across hematologic malignancies. The location of pre-B-ALL and E/R+ samples are indicated on the plot. Red color tones indicate high expression. c. Distributions of expression level at diagnosis and day 15 post-treatment are shown as ridge plots for a set of TFs with significant expression change (Table S2). X-axis corresponds to normalized expression level. The differentiation marker *MS4A1* (mRNA) / CD20 (corresponding protein) level is shown for comparison. d. Candidate regulatory interactions evaluated at active enhancer and promoter regions (highlighted by shading) at the *ELK3* locus. TF regulons that included *ELK3* with nPred>4 are shown. The ChIP-seq peaks are shown at these locations, corresponding to HUVEC (ELK3), HSC (ERG, FLI1, GATA2), K562 (IRF1 upon IFNy stimulus) and GM12878 (MEF2C and RFX5) peak annotations (no data for ZNF467 available). Lines between enhancer or promoter regions and regulons are drawn when corresponding peak was found. GRO-seq data is shown from E/R+ REH cell line and two primary E/R+ bone marrows. e. The luminescence signal from MTS assay (above) and relative cell counts (viable cells in colored bars, total cell count indicated without fill) at different drug concentrations are shown TK: TK216, XR: XRP44X).

In further confirmation, we analyzed TF expression matching the positive TF regulons with high activity in E/R+ cells (top panel, Fig. 5a) across a large microarray gene expression dataset [50] (Hemap, N=9544, with 1304 pre-B-ALL samples). The ETS-factor *ELK3* and *SP4* have been implicated by genome-wide association (GWAS) studies as risk loci for pediatric pre-B-ALL [48,49]. Based on the bulk transcriptomes, we could validated both to be expressed in B-ALL, with highest proportion detected in the E/R+ subtype (red arrow), as shown comparing hematologic malignancies on the t-SNE plot of Hemap samples (Fig. 5b), where lymphoid malignancies are highlighted above the panel (Fig S6). The two most common B-ALL subtypes (E/R+ and high hyperdiploid cases) displayed similarly high *ELK3* and *ERG* expression (Fig S6). Overall, we could confirm the expression in E/R+ leukemias (log2 signal above probe detection level of approximately 6 (Fig S6)) for all 19 TFs analyzed.

### Leukemic TF activities that persist during chemotherapy provide new targets to overcome resistance

Next, we analyzed the effect of the standard leukemia induction therapy (prednisolone, vincristine, doxorubicin) on the TF expression based on the scRNA-seq profiles acquired at mid-induction therapy in ALL10 and ALL12 (Fig. 5c). Based on differential distribution analysis, residual leukemic blasts from day 15 bone marrow had lower expression of *RUNX1, TCF3, SOX4* and *ERG* compared to diagnostic state in both samples, while *SMAD1* and *ELK3* levels increased slightly (refer to Table S2 for full analysis). ALL10 had a favorable decrease in blast count at end of induction on day 29 (0.08%). At day 15, the expression of pre-/immature-B TFs *POU2F2, KLF2/6, AFF3* and *SPIB* were elevated in the remaining leukemic cells of ALL10 (10% blasts). These changes may relate to the differentiation-inducing effects of glucocorticoids (daily prednisolone). However, overall the changes in TF activities or gene expression were modest, indicating that only partial differentiation towards pre-B cell state may occur, despite the increase in the maturation marker CD20 (encoded by *MS4A1).* In contrast, cases ALL3 and ALL12 responded slowly to therapy (74% and 59% blasts at day 15; 0.16% and 0.2% end of induction, respectively). In ALL3, the cell cycle state distribution was strongly skewed to G0/G1 state at diagnosis (Fig. 3d) compared to the other E/R+ cases, which could underlie resistance to drugs targeting dividing cells (doxorubicin/vincristine). In ALL12, the day 15 sample TF profile indicated persistence of the leukemic gene regulatory program, manifest as lack of pre-/immature-B TF upregulation (Fig. 5c).

This prompted further analysis of TF regulation in E/R+ cells. Towards this end, we extracted predicted regulatory interactions for *ELK3* from TF regulons (Npred>4) and analyzed the active enhancers and transcripts based on global run-on sequencing (GRO-seq) data (Fig. 5d). The GRO-seq profile confirmed high transcriptional activity of the gene locus in E/R+ cells (E/R+ cell line REH and two primary E/R+ bone marrow profiles are shown) and revealed an alternative TSS upstream the annotated *ELK3* TSS. We retrieved public ChIP-seq peak data for the TFs that could represent *ELK3* upstream factors, and narrowed down the analysis to active promoters and enhancers detected by GRO-seq: ELK3 (data from HUVEC) and the two other ETS-factors ERG, FLI1, together with GATA2 (data from HSC) bound the novel TSS, while ELK3-FLI1-GATA2 or FLI1-ERG-MEF2C bound at intronic enhancers (Fig. 5d). Further, their binding may contribute to the aberrant cytokine, chemokine and growth factor expression detected in E/R+ cells based on regulon target and ChIP-seq peak data (Fig. S6a). Thus, the regulon and ChIP-seq analysis indicates that a tightly interconnected network may form between TFs with high activity in E/R+ cells.

As one strategy to overcome resistance to standard induction therapy, we sought to identify drugs that could target the high activity TFs identified. We selected two compounds for further experiments: XRP44X has been shown to have dual activity in both targeting microtubules (like vincristine) and simultaneously decreasing ELK3 activation by inhibiting its phosphorylation [51]. TK216 is an analog of YK-4-279 that was directly shown to inhibit ERG and FLI1-mediated transcriptional activity [52]. We used the glucocorticoid-resistant E/R+ REH cells as a cellular model and performed proliferation and viability assays at different drug doses (Fig. 5f; Fig. S6b. At 72 h, cellular ATP levels assessed using MTS assay (top panel) and viable cell counts (bottom panel) dropped sharply at sub-micromolar doses of XRP44X and TK216. Moreover, >1 uM doses (1.6 uM for XRP44X, 2 uM for TK216) resulted in loss of cellular ATP. In summary, small molecule inhibitors targeting the leukemic regulatory network could be effective in drug-resistant leukemic cells.

## Discussion

Specific cell types are faithfully generated in a repeated manner during development. This is due to gene regulatory interactions that limit the space of stable cell states [53]. Understanding the direct impact of aberrant leukemic TFs on cell state transitions in differentiating lymphoid cells, and identifying such TFs that maintain leukemia-specific cell states could enable more precise therapeutic intervention. Here, we explored large-scale single cell transcriptomics data from healthy human BM to generate a reference for cell state transitions and TF activities that characterize early B-lineage differentiation. Focusing on leukemias carrying the E/R fusion, we profiled primary patient bone marrow samples from diagnosis and during induction therapy. The data suggest that the leukemic cell states resemble most the pro-B state, differ between cases in their cell cycle activity, interact with the immune microenvironment and may partially be programmed towards pre-B state during chemotherapy. Accompanying the differentiation arrest at pro-B cells, our results revealed elevated activity of specific TFs that could serve as therapeutic targets.

Single cell profiling techniques have challenged how we define cell types and provided new methodology to characterize their molecular phenotypes [23,54]. Previous analysis of the HCA BM data [2,27] distinguished the B-lineage cell populations but did not further compare them, or analyze how the transition from HSC to immature B cells is regulated. One distinguishable feature along this lineage are the alternating cycling and G1 cell populations that the single cell profiling uniquely could resolve. Here, we focused on uncovering key lymphoid TFs orchestrating these cell state transitions. A popular approach to study gene regulation based on scRNA-seq profiles is to analyze so-called TF regulons defined by TF-to-target correlation and TF motif analysis, available in the SCENIC tool [24]. We benchmarked this method for studying BM cell states, using target genes for EBF1, PAX5 and BCL11A from ChIP-seq as validation. Compared to the original method, we introduced a cross-validation step and improved capture of repressive TF-target interactions. The regulons thus identified faithfully captured targets confirmed by ChIP-seq and TFs that have been previously functionally implicated in B-lineage differentiation through mouse knockout studies [32,33]. This same analysis strategy could be adopted to identify candidate regulatory programs for cell states across hematologic malignancies.

In this study, we examined the TF activities that may contribute in maintaining leukemic cell states in E/R+ cases and linked those to target genes, including modulators of leukemia-immune cross-talk. Previous bulk cancer genomics studies have established that repeated gene expression patterns also characterize cancer samples [55], including ALL where such studies have established several transcriptome-based subtypes [56–60]. They have also shed light on pathway activity and TF expression in E/R+ cells that could be utilized to design targeted therapies [6,61,62]. However, the rarity of normal B-lymphoid pro-B cells in BM tissue has represented a challenge to perform direct comparison of E/R+ and healthy BM lymphoid cell states *in vivo.* Moreover, bulk profiles have obscured the characteristics of the immune microenvironment. Existing scRNA-seq studies in ALL have so far not focused on the these aspects, in particular on the leukemic gene regulatory network [63,64]. Through computational discovery and analysis of TF regulons from scRNA-seq data, and independent validation with GRO-seq and bulk genomics data, we could show that elevated activity of multiple ETS-factors (ELK3, ERG and FLI1), together with pro-B TFs FOXO1, MEF2C, immature B-cell TFs NFYC, RFX5, lineage-atypical GATA2 expression and E/R-subtype-specific SP4 and TCFL5 activities characterized the E/R+ regulatory network. *TCFL5* has been previously shown to be upregulated in E/R+ pre-B-ALL [65–67], while GATA2 has been reported to contribute in the upregulation of erythroid genes, such as *EPOR*, a known marker gene in E/R+ leukemia [68–70]. While these TF activities were consistently high across the six diagnostic samples studied, many IRF- and STAT-regulons showed variable activity. Previously, inhibition of STAT3 was tested in E/R+ leukemic cells and shown to be necessary for *MYC* expression [61]. However, we did not observe the correlation between STAT3 and MYC regulon activities in our analysis.

Among the E/R+ TF network, *ELK3* and *SP4* have been reported to confer risk of leukemia development in GWAS studies [48,49]. Previous expression quantitative trait loci data from mature B lymphoid cells indicated that the *ELK3* risk variant associates with its lower expression [48]. This contrasts the data obtained here where high expression was seen in E/R+ scRNA-seq data, which we confirmed by bulk gene expression data comparing across hematologic malignancies [50]. This data indicated similar expression levels also in high hyperdiploid pre-B-ALL samples that represent the most common ALL subtype. In E/R+ cells, we observed an active alternative TSS at the *ELK3* locus. Further functional studies on the impact of the risk variants on expression of *ELK3* variants in normal pro-B cells and leukemia are thus warranted to characterize their role during leukemogenesis. One aspect to study in this context is the role of immune surveillance of pre-leukemic clones, as the target genes that were reproducible associated with the ELK3 regulon across SCENIC runs included *TGFB1, TERF2* and *HLA-E* that we showed to be highly expressed in E/R+ cells. In addition to HLA-E, class I MHC molecules HLA-A, -B, C, and F were also upregulated in leukemic cells. Functionally, their expression might interfere with NK cell-mediated tumor surveillance [40,42–44,66,71,72]. It is known that infection exposure is a key underlying factor in the development of E/R+ leukemias [73–76]. The decrease in NK cell number observed in the five E/R+ BM characterized here is in agreement with a larger flow-cytometry based study [41]. However, using scRNA-seq data from E/R+ and normal BM, we could analyze the small NK cell population further. There was a shift towards immature NK cell populations in leukemic BM and we did not detect subpopulations with high BHLHE40 or IFNγ expression that would characterize active tumor killing, matching targets inhibited by TGF-β [77,78]. Interestingly, the TF regulons did not indicate canonical activation of SMAD2/3 by TGF-β in the E/R-leukemic cells but instead both the regulon and differential expression analysis showed high *SMAD1* levels. Atypical activation of SMAD1 via TGF-β has been reported to occur in different cell types [79,80], and instead of suppressive signaling it may give E/R+ preleukemic cells a growth advantage over healthy pro-B cells [73]. Overall, our analysis provides a rationale for carrying out further studies focused on immune cell-leukemia cross-talk to develop therapies that specifically target these immune cell suppressive mechanisms and to understand how genetic variations in the leukemia-associated TF loci relate to leukemia risk.

Measurable residual disease (MRD) at mid-[90] and end of induction chemotherapy are predictive markers for relapse risk [13]. Moreover, *in vitro* resistance to prednisolone has been shown to confer poor prognosis [91]. Previous bulk gene expression studies have indicated treatment-specific changes in gene expression and expression of more mature cell markers [92,93]. In this study, we sought to gain insight on the efficacy of drug therapy in leukemic cell clearance examining cell state features from scRNA-seq samples collected during *in vivo* chemotherapy. The E/R+ samples analyzed included several cases with residual leukemia cells at mid (day 15) or end of induction (day 29), and we profiled two of these from post-therapy bone marrow at day 15. ALL10 with a favorable end of induction blast count (<0.1%), regained expression of multiple pre-B/immature B-specific TFs, including *SPIB* and *AFF3.* In contrast, similar changes in TF expression were lacking in blasts representing 59% of BM cells in ALL12 at day 15. In ALL3 that had also a high blast count at day 15, the leukemic blasts at diagnosis represented predominantly non-cycling cells. Characterization of these features across a larger patient cohort is thus warranted. To overcome resistance to standard induction therapy, our analysis highlighted candidate drug therapy targets in E/R+ cells that could disrupt leukemic TF activities. We tested small molecule drugs targeting the ETS-factors ELK3, or ERG and FLI1 in dexamethasone resistant E/R+ REH cells and found reduced cell viability with sub-micromolar concentration. Inhibitors abrogating FLI1, MEF2C, ELK3, or SP4 activation have been previously shown to have efficacy in different cancers [94–99]. Moreover, the small molecule ERG/FLI1 inhibitor tested here has entered a phase 1 study in Ewing sarcoma [100]. Our findings demonstrate the feasibility of monitoring the early treatment response using single cell genomics characterization and its potential to uncover targets for further pre-clinical and clinical studies.

One limitation of this study is that we could only compare the E/R+ leukemic cells to early B-lineage differentiation in adult BM. In our analysis, a putative steady state of pro-B cells in G1 state was connected to the succession of cell states from early lymphoid to pre-B state. Pro-B cells can migrate during early development from fetal liver and contribute as a progenitor pool to lymphoid cell generation alongside HSC during early life [34]. As pre-leukemic clones may arise already *in utero*, the origin and the relative contributions of both HSC- and pro-B pool-derived lymphoid cells at different ages would be relevant to characterize further, which could be achieved using new lineage tracing approaches coupled with scRNA-seq [101–103]. Moreover, compared to other hematopoietic lineages, the succession of lymphoid cell states from early lymphoid to immature B cells differed markedly in transcriptional activity and cell size. The sequential transitions between G1 and cycling cell states pose challenges in single cell analysis in data normalization and resolving the B-lineage differentiation path. Existing benchmarks with down-sampling of counts [104,105] show that normalization methods are robust to differences up to 20% in “size”, yet the differences between G1 and G2/M states observed in lymphoid cell data exceeded this. Moreover, many common trajectory analysis methods fit tree-like structures to data [106]. This challenge motivated our choice of diffusion pseudotime and RNA velocity analyses that both can accommodate cycling cell states[25,31,107]. The variability between donors in relative proportions of cycling cells at each differentiation state, would also represent a confounder in comparative analysis of cells categorized using differentiation markers alone, as carried out in previous flow sorted bulk transcriptomes. Therefore the comparisons of subsequent differentiation states matched by cell cycle state, as performed here, represents a significant advance. One technical confounder in scRNA-seq performed using viably frozen (unfixed) BM samples could derive from the specific protocol used for thawing cells, which could introduce differences in the transcriptional activity level of cells measured. We noted that the largest variance (PC1) within individual leukemic bone marrow samples reflected their transcriptional activity. These effects could be mitigated by careful selection of analysis steps and underline the importance of good benchmarking data for optimizing single cell workflows for clinical samples.

## Conclusions

This study provides the first comprehensive characterization of cell states and TF activities in E/R+ ALL cases and its comparison to normal human B-lineage differentiation at single cell resolution. Through joint analysis of single cell and bulk genomics data, we characterized TF activities contributing to the aberrant cell phenotype in leukemic cells. These results could provide a rational basis for developing new therapies targeting leukemia-immune cell cross-talk and treatment-resistant leukemic cell states.

## Methods

### Patient samples

This study was approved by the Regional Ethics Committee in Pirkanmaa, Tampere, Finland (#R13109) and conducted according to the guidelines of the Declaration of Helsinki. A written informed consent was received by the patient and/or guardians. All the patients were positive for the E/R-fusion transcript based on clinical RT-qPCR and FISH analysis (further confirmed using bulk WGS data). Their age ranged between 1-10 years and all cases received standard induction therapy according to the NOPHO ALL-2008 protocol, with prednisolone 60 mg/m^2^/day p.o. days 1-29, vincristine 2.0 mg/m^2^ i.v. days 1, 8, 15, 22, and 29, doxorubicin 40 mg/m^2^ i.v. days 1 and 22, and methotrexate i.t. days 1, 8, 15, and 29 [108]. Leukemic blast percentages in the bone marrow during diagnosis, at day 15, and at day 29 are shown in Table 1. Mononuclear cells (MNC) were extracted from fresh BM using Ficoll-Paque Plus (GE Healthcare, #17-1440-02). Bone marrow MNCs were also extracted from two patients (ALL10 and ALL12) during the induction therapy at day 15 after initiation of therapy. MNCs were viably frozen in 15% DMSO/40% FBS in RPMI in liquid nitrogen. In addition, nuclei were isolated for GRO-seq (ALL7 and ALL13) as described in [5], snap-frozen and stored at −80**L** in a freezing buffer containing 40% glycerol.

### Cell line samples

The E/R+ REH cell line (ACC-22, DSMZ, Germany) was maintained in RPMI 1640 (Gibco, Thermo Fisher) supplemented with 10% FBS (Gibco, Thermo Fisher), 2 mM L-glutamine (Gibco, Thermo Fisher), penicillin (100 U/ml), and streptomycin (100 mg/ml) (Sigma-Aldrich). Mycoplasma status was defined negative for all cell lines by PCR (PCR Mycoplasma Test Kit I/C, PromoCell GmbH, Germany) and cell lines were authenticated by Short Tandem Repeat genotyping (Eurofins Genomics, Ebersberg, Germany).

### scRNA-seq

Single cell gene expression was studied to characterize leukemic bone marrow cell populations. Cells from primary BM samples (n=6 diagnostic, n=2 post-treatment) were processed for scRNA-seq in the Finnish Functional Genomics Center, Turku, Finland, in 4 batches: 1) ALL3, 2) ALL1 3) ALL10 and ALL10-d15, and 4) ALL8, ALL9, ALL12, and ALL12-d15. Before applying the cells into the Chromium cartridge, their viability was checked using Trypan blue. PI-negative (live) cells were selected from sample ALL3 using FACS. Samples ALL1, ALL10, and ALL10-d15 were processed directly after thawing the MNC fraction without further processing. Excess dead cells were depleted from samples ALL8 and ALL9 using bead-based Dead Cell Removal Kit (#130-090-101, MACS miltenyi Biotech), increasing the percentage of viable cells from 43 % to 72 % and from 63 % to 78 %, respectively. For samples ALL12 and ALL12-d15, enrichment of leukemic cells was carried out by depleting non-B-cells using streptavidin-beads (BD Streptavidin Particles Plus, BD Biosciences, Franklin Lakes, NJ, USA) and biotinylated antibodies against human CD16 (clone 3G8), CD14 (HCD-14), CD11c (3.9), CD56 (HCD56), CD3 (UCHT1), and CD66 (G10F5) (Biolegend), all with final concentrations of 2 ug/ml, following the manufacturer’s instructions and as previously described (Good et al. 2018 Nat Meth). Depletion efficiency was estimated by flow cytometry using CD3 (BV421, BD Biosciences, #56287, RRID:AB_27378607) and CD19 (Thermo Fisher Scientific, # 25-0199-41, RRID:AB_1582279) antibodies, with a viability dye (eBioscience, Fixable Viability Dye eFluor™ 506, #65-0866-14). Depletion decreased the proportion of T-cells (CD3+) from 30 to 2%, increased the proportion of B-cells (CD19+) from 23 to 50%, and increased the percentage of viable cells from 50 to 80% in a test BM sample.

scRNA-seq was performed using the 10X Genomics Chromium technology, according to the Chromium Single-Cell 3’ Reagent Kits V2 User guide Rev B. In brief, cells were combined with reverse transcriptase Master Mix and partitioned into Gel Bead-In EMulsions (GEMs) using 10X GemCode Technology, where the poly-A transcripts are barcoded with an Illumina R1 sequence, a 16 bp 10X barcode and a 10 bp Unique Molecular Identifier (UMI). 11-12 cycles of PCR was used to amplify the cDNA. Sequencing was performed using the Illumina HiSeq 3000. Primary BM samples were sequenced to an average depth of ~50 000 reads per cell.

### HCA bone marrow scRNA-seq data processing and cell state annotation

Characterization of normal bone marrow B-lymphoid cell states was performed using data from healthy donors (n=8), available from the HCA data portal. Raw fastq-files corresponding to 10X Genomics Chromium single cell data were downloaded from [109]. Data was aligned with Cell Ranger 3.0.2 to human reference (hg19) version 3.0.0 with default parameters and the filtered count matrix was taken for downstream analysis. Scanpy [110] (version 1.4) was used for initial characterization of cells[111] as follows: Genes were first filtered to include only genes present in more than 100 cells. Then, bad quality cells were removed if i) UMIs arising from mitochondrial genes in a cell accounted for more than 10% of total UMI count, while possible doublets were excluded based on ii) total number of UMIs 50 000 or more, or iii) the number of genes expressed in a cell 6 000 or more. Next, genes were filtered once more to include only those expressed in minimum 400 cells. Highly variable genes (HVG) were defined as genes with minimum mean expression 0.0125, maximum mean expression 3, and minimum dispersion 0.5, resulting in 2046 genes with the rest of the genes filtered out from the data for downstream analyses. To reduce undesired technical effects in data analysis, the function “regress_out” was used to normalize the effect of the number of UMIs and the percentage of UMIs arising from mitochondrial genes to gene expression in each cell. Mutual nearest neighbors (MNN) correction [112,113] (mnnpy version 0.1.9.5) was used to combine data across the eight donors for clustering and cell state identification (batch_categories set to Manton donor identifier). Principal Component Analysis (PCA) was then calculated using the processed data (Scanpy version 1.4). Top 50 principal components were used to calculate a neighborhood graph (number of neighbors was set to 30) that was used as input for Uniform Manifold Approximation and Projection (UMAP) [114], where min_dist was set to 0.5, and Louvain clustering [115] with resolution set to 1.0, which was enough to characterize major cell type clusters from the data. Wilcoxon test was used to find marker genes for each cluster which were used to characterize the found clusters in concordance with known marker genes. Cell cycle states (G1, S, G2/M) of cells were annotated using score_genes_cell_cycle-function in Scanpy using annotated cell cycle genes from [116].

To focus on B-lineage cell differentiation, a subset of cells from clusters containing hematopoietic stem cells and B cell lineage cells was re-analyzed in an iterative manner, each time running the basic workflow again with additional filtering steps. Initially, genes expressed in less than 100 cells were removed when analyzing this subset. When choosing highly variable genes, we required the minimum dispersion to be 1, compared to the previous 0.5. Small clusters containing high expression of markers for T cells, NK T cells, monocytes and erythroid precursor cells were still present after the first iteration and were filtered out. In the second iteration, we required the minimum mean expression to be 0.1 and the minimum dispersion 0.5 for choosing highly variable genes. In the neighborhood graph calculation, the number of principal components used was set to 20. Next, we filtered each cluster for possible outliers by calculating cluster-specific Absolute Median Deviance (MAD) for number of UMIs and percentage of UMIs from mitochondrial genes and removed cells assigned to the cluster with MAD greater than 5 in either. This was motivated by the large differences between clusters in these metrics. During B cell differentiation, the cells display marked changes in cell size (e.g. transitioning from large cycling pre-B cells to small pre-B cells). Thus this choice is also motivated by biology. With the filtered subset of 20 753 cells we ran through the workflow once again, choosing highly variable genes with minimum mean 0.1 and minimum dispersion 0.75 and set the number of principal components in neighborhood graph calculation to 20. The final clusters were characterized as described above.

### ALL scRNA-seq data processing and cell state annotation

To perform similar analysis in leukemic BM, raw patient data acquired in this study (n=6 diagnostic, n=2 post-treatment) was processed and aligned with Cell Ranger (version 3.0.2) with the same settings as the HCA data. Scanpy (version 1.4) was used for initial characterization of cells following the same approach as outlined above [111] (HCA analysis): Genes were first filtered to include only genes present in more than 100 cells, requiring this metric to exceed 200 in the final iteration. Cells were removed if i) UMIs arising from mitochondrial genes in a cell was more than 10 %, ii) total number of UMIs was 40 000 or more, or iii) number of genes expressed in a cell was 5 000 or more. Highly variable genes were defined as genes with minimum mean expression 0.0125, maximum mean expression 3, and minimum dispersion 0.5, resulting in 1425 genes that were used for clustering and dimensionality reduction (50 principal components, number of neighbors 15, resolution 1.0). MAD filtering was used to remove outlier cells from clusters, as described above. With the final cell subset passing these criteria (44746 cells), the workflow was repeated and clusters characterized based on marker genes.

### Differential distribution of read counts: scDD analysis

The gene expression distributions in subsequent cell states representing B-lineage differentiation, or between leukemic and normal cell states, were analysed with the scDD-package [26,111]. The tool enables comparisons based on differential distribution and proportion of zeros between two groups of cells. Genes were assigned into three main categories - DE, DM, and DZ. DM and DE characterize changes in the expression distribution in cells with non-zero count for the gene analyzed (differential mean and differential modality, respectively). DZ genes differ between the groups in proportion of cells with zero read count for the gene analyzed. In the context of differentiation, where cells switch genes on/off to proceed in maturation, this metric captured majority of the changes.

To account for differences in the number of UMIs and genes detected in different cell types, variance stabilizing transformation [105] (version 0.2.0) was used to correct for these differences before differential distribution testing. Sample was used as the batch interaction term and logarithm of UMI counts in cell (“log_umi”) was specified as the latent variable to regress out. The resulting corrected UMI counts were then used as input to scDD. When running scDD we noticed that for some genes the clustering of the expression level within scDD failed due to zero variance. To overcome this, the scDD tool was modified to add a small jitter to genes which had this problem [117]. Cells with 3000-3500 counts after the corrections were included in comparing the pre-B G1 vs. pro-B G1, and the pro-B G1 vs. leukemic G1 cells. The following number of cells per differentiation/disease state were compared: HSC 3660; early B-lymphoid 895; pro-B cycling 794; pro-B G1 1413; pre-B cycling 1714; pre-B I G1 2541; pre-B II G1 2025; diagnostic leukemic G1 6340; diagnostic leukemic cycling: 7054.

Further filtering for scDD results was done using adjusted p-value and fold change or difference in percentage cut-offs (Fig S2). P-values were adjusted using the Benjamini-Hochberg FDR method.

### Clustering genes based on differential zero proportion

Differentially distributed genes from the leukemic vs. pro-B zero proportion comparisons, present in both G1 and cycling cell based comparisons (90 downregulated and 272 upregulated), were clustered based on their zero proportion metric in ten cell states (HSC, early lymphoid progenitors, proB cycling (S/G2/M), pro-B G1, pre-B cycling, pre-B G1 I, pre-B G1 II, immature B, leukemic cells G1, and leukemic cells cycling). K-means centroids were calculated using the R package flexclust [118] (version 1.4-0) with k = 8 and correlation as distance metric using the kccaFamily function (cent = centMean, dist = distCor).

### Pathway enrichment analysis

Gene lists were analyzed for enrichment of ontology and pathway terms using the online web server Enrichr [119,120] (release Jan 2019). The analysis was performed based on gene sets from GO, MGI Mammalian Phenotype, Reactome and TF perturbations. Enriched terms were selected based on the combined score (>150) cut-off. The combined score refers to the combination of p-value (Fisher’s exact test) and the z-score that represents the deviation from the expected rank.

### Ordering cells based on pseudotime

Pseudotime analysis can be used to find a latent trajectory (pseudotemporal ordering of cells) in single cell data, corresponding to differentiation or cell cycle. HSC and B lineage cells from HCA BM data were subjected to pseudotime analysis following the best practices workflow by Luecken and Theis [121] using Scanpy (version 1.4.5). Non-expressed genes (zero UMIs in any cell) were excluded and the data was normalized with size factors calculated with computeSumFactors-function from scran-package [104,122] (version 1.10.2) where louvain clusters (resolution 0.5) were used and min.mean was set to 1. The analysis was done two ways: using highly variable genes or selecting differentially distributed genes from our scDD analyses between HSC and B lineage cell types and the cell cycle phase marker genes.

Neighborhood graph was calculated with the number of principal components set to 15 and the number of neighbors set to 15. Diffusion map representation [123] was then calculated obtaining 15 diffusion components and a pseudotime ordering was calculated using diffusion pseudotime [107] using 10 diffusion components and setting the required root cell as the HSC with the highest value in the 1st diffusion component (DC1). For visualization the DC1 vector was mirrored to obtain a left to right pseudotime trajectory of cells. The ordering of clusters was highly comparable with HVG or custom gene selection. The latter is shown in figures for consistency.

### RNA dynamics analysis

During differentiation, dynamic changes occur in gene transcription that can be modelled based on newly synthetized RNA (reads corresponding to unspliced mRNA) and processed RNA (reads corresponding to mRNA). Based on the dynamic RNA processing model, predictions of the future transcriptome state can be obtained and visualized together with the measured current state. Velocyto CLI [25] (version 0.17.17) was used to calculate spliced and unspliced counts per gene using human reference genome (hg19) version 3.0.0 for Cell Ranger from 10x Genomics. Expressed repetitive elements were masked using expressed repeat annotation for hg19 downloaded from UCSC Genome Browser [124]. scVelo-package [31] (version 0.1.21) was used to analyze RNA dynamics in B cell differentiation. The gene expression matrix was accompanied with the spliced and unspliced count matrices of HSCs and B lineage cells from HCA BM data. The data was first filtered by removing genes with less than 10 shared counts in both spliced and unspliced data. The matrices were each then normalized by dividing the counts in each cell with the median of total counts per cell. The 3000 most variable genes were extracted based on the spliced count matrix and the data matrices were log-transformed. 30 top PCs were defined based on the most variable gene spliced count data followed by neighborhood graph calculation, with the number of neighbors set to 30. Based on the neighborhood connectivities, the first order moments for spliced and unspliced matrices were calculated. The normalized unspliced and spliced count matrices were then used to estimate the velocity of each cell using the deterministic model. The velocities were embedded to a UMAP presentation which was calculated with the same preprocessing steps before calculating the diffusion map.

### Regulon discovery and TF activity scoring

For the discovery of TF activities that characterize specific cell states, a modified SCENIC workflow [24,111] was developed based on the python implementation of the SCENIC method [125]. In our implementation, equal amounts of cells per cell type were sampled from the original data to ascertain that differences in cell type abundances do not bias the analysis. Secondly, the discovered regulons were evaluated based on a left-out test set. Specifically, the input matrix (equal representation of cell types) was split into training (70% of cells) and test (30% of cells) sets. The default SCENIC pipeline for regulon discovery was then run for the training set. The regulons found were scored in the training and test sets and and the average score per cell type calculated in both sets. These mean regulon scores across cell types were compared between training and test sets with Pearson’s product moment correlation coefficient. Regulons with p-value > 0.001 were discarded. The discovery was repeated 10 times. The final set of regulons was then scored using the whole original data set. Because different iterations often find regulons with the same driving TF and a similar target gene set, the mean score of the regulon for each cell was used in downstream analysis. In these analyses, leukemic cells from different donors and collection times were treated as separate cell types. For filtering regulons, a linear model was fit 100 times to a random subset of the regulon score matrix where 600 cells per cell type were sampled from the original data set. In the model, the response is the regulon score and the cell type label is the independent variable (score ~ cell type). Regulons with the coefficient of determination (R^2^) < 0.5 were considered to not show sufficient variation between cell types and were therefore filtered out (Table S3 shows the unfiltered result). Additionally, a regulon was filtered out if the mean score in any cell type was above 70% percentile while the TF’s gene expression had > 95% of zeros.

### Cell type assignment of ALL cells with label transfer

Annotated HCA BM cells were used as a reference to label the ALL scRNA-seq data. This was performed with label transfer functions from Seurat [39] (version 3.1.4) as follows: Each ALL sample was separately normalized with CPM with scale factor of 10000 and then log-transformed followed by extracting top 2000 most variable genes. Then, separately for each ALL sample, transfer anchors between reference and sample were calculated with FindTransferAnchors-function where the first 30 dimensions of CCA were used as neighbor search space (parameter *dims).* Finally, the TransferData-function was used to annotate the leukemic cells with 30 first PCs used in the weighting procedure.

### NK cell analysis

Clusters labelled NK and NK T cells from full HCA BM and primary ALL data were combined and processed together starting from raw counts with Scanpy (version 1.4.5). Genes were first filtered to include only genes present in more than 100 cells. Then cells were removed if i) UMIs arising from mitochondrial genes in a cell was more than 5 %, ii) total number of UMIs was below 500 or 3 000 or more, or iii) number of genes expressed in a cell was below 200 or 3 000 or more. Then, data was normalized with following the same steps and parameters as in the pseudotime analysis followed by extraction of 3 000 most variable genes which were used to calculate the first 50 PCs followed by neighborhood graph calculation with the 50 PCs and n_neighbors set to 15. Leiden clustering [126] with resolution 1 was calculated identifying two clusters with high expression of erythroid markers *HBA1, HBA2* and *HBB* which were then removed and analysis repeated starting from calculating the most variable genes. UMAP embedding was calculated with the obtained PCs and the neighborhood graph to visualize the data. Leiden clustering was calculated again but with resolution parameter set to 2 to obtain more detailed clusters. NK clusters were separated from NK T clusters based on *CD3D* expression. Marker genes for the clusters were calculated with the rank_genes_groups-function with method-parameter set to “t-test_overestim_var” (Welch t-test) and discarding genes with fold change less than 2. Top 5 genes per cluster based on test score were extracted. Scores for NK subtype gene sets from [45] were calculated with the score_genes-function using the top 20 genes per gene set sorted by log-fold change.

### Bulk pro-B cell ATAC-seq analysis

For analyzing open chromatin regions in pro-B cells, ATAC-sequencing data of human fetal pro-B cells (n=3) were retrieved from NCBI SRA database, GSE122989 [127]. Data pre-processing and peak calling was done following the ENCODE pipeline for ATAC-seq [128] (version 1.5.4) which is a tool for statistical signal processing and produces alignment and measures of enrichment. Caper configuration file was set up for the local server platform and parameters in the JSON file were selected based on the example JSON file. Hg19 was used as a reference genome in alignment. Narrow peaks were pooled and merged from three replicates. The highest enriched 10 000 peaks were taken to downstream analysis. Regions overlapping annotated TSS (NCBI RefSeq and UCSC Known gene) were discarded. TF motif discovery was performed with HOMER [129] (version 4.9.1) findMotifsGenome.pl (-size 200 -mask) using the remaining (3923) open chromatin regions. P-values were adjusted using the Benjamini-Hochberg FDR method.

### GRO-seq assay

To study enhancer and gene region activity, primary ALL BM samples (n=2) were collected for GRO-seq. In addition, our existing data in REH cells available via NCBI GEO (GSE67540 [132]) were analyzed. For these samples and ALL7, the nuclear isolation and library preparation protocols were performed as described in [12]. Briefly, run-on products labelled with BrUTP were extracted with TRIzure (Bioline, London, UK). RNA was precipitated first for 30 min on RT and then for an extra 10 min on ice. Poly-A tailing reaction was carried out and nascent RNA collected using anti-BrUTP beads. The anti-BrUTP beads used previously [12] were not available for the collection of run-on products for ALL13 and for this sample the libraries were performed as described in [133] with few modifications. Bead-binding was performed using 30 μl of Protein G Dynabeads (Thermo Fisher Scientific Baltics UAB, V.A. Graiciuno 8, LT-02241 Vilnius, Lithuania. Thermo Fisher Scientific Baltics UAB complies with Quality System Standards ISO 9001 and ISO: 13485) per sample with 2 μg anti-BrdU monoclonal antibody (cat# ab6326, Abcam, Cambridge, UK). Beads were washed four times with 300 μl of PBST wash buffer including SUPERase In RNase Inhibitor (Thermo Fisher, Carlsbad, CA, USA). The purified run-on RNAs were next converted to cDNA and PCR amplified for 13 cycles size and selected to 225-350 bp length. Single-end sequencing (50 bp) was performed with Illumina Hi-Seq2000 (GeneCore, EMBL Heidelberg, Germany).

### GRO- and ChIP-seq data pre-processing

TF ChIP-seq was used to validate TF-target associations obtained using SCENIC. ChIP-seq data representing PAX5 and EBF1 (GSE126300 [137]) were available in hg19, while BCL11A (GSE99019 [138]) read data was processed to hg19 from raw reads. For BCL11A and GRO-seq data, the raw sequencing reads were quality controlled using the FastQC tool [134]. Bases with poor quality scores were trimmed (min 97% of positions have a min phred quality score of 10) using the FastX toolkit [135]. Duplicate reads were collapsed from ChIP-seq files using fastx (collapse), while reads mapping to rRNA regions (AbundantSequences as annotated by iGenomes) were discarded from GRO-seq data. The Bowtie software [136] (version 0.12.9 for GRO-seq, version 1.2.3 for ChIP-seq) was then used for alignment of remaining reads to the hg19 genome version, allowing up to two mismatches and no more than three matching locations. The best alignment was reported. Reads overlapping with so-called blacklisted regions that include unusual low or high mappability as defined by ENCODE, ribosomal and small nucleolar RNA (snoRNA) loci from ENCODE and a custom collection of unusually high signal depth regions from GRO-seq was used to filter the data. Subsequently, data was analyzed using HOMER [129] (version 4.9.1). GRO-seq tagDirectories were generated with fragment length set to 75 and data visualized using makeMultiWigHub.pl with strand-specificity. HOMER [129] (version 4.9.1) findPeaks tool (-style factor) was used in peak calling from ChIP-seq against input sample.

### ChlP-seq peak analysis

The peak data was ranked based on peak calling statistics (lowest rand corresponding to best peak) and the rank annotated in each peak name. Next, peaks were associated with nearby genes using the approach described in [35]. The data was summarized by gene, recording the number of associated peaks, the peak ranks and peak distances to gene TSS.

### Immunofluorescence stainings and flow cytometry

For flow cytometric analysis, 500 000 cells were washed with Cell staining buffer (catalog number #420201; all from Biolegend, San Diego, CA, USA, if not stated otherwise), and FcR was blocked using TruStain human FcX for 5 min (#422301). For intracellular staining, 150 μl of Cyto-Fast Fix/Permeabilization buffer (#426803) was added to 100 μl of cells in cell staining buffer, and incubated for 20 min at room temperature in the dark. Cells were washed 2X with Cyto-Fast Perm Wash solution and stained with antibodies and kept on ice in the dark for 20 min during staining. All centrifugations were done at 350 x g. Cells were washed 2X with Cell staining buffer and measured with Accuri C6 (BD Biosciences, CA, USA). RNA flow analysis was performed with Amnis® FlowSight® imaging flow cytometer (Luminex Corporation, TX, USA).

### Cell proliferation and viability

Effect of drugs targeting TF activities that were found to be high in E/R+ leukemia was studied in the glucocorticoid-resistant REH cell line. The experiments were performed in three biological replicates. TK216 (ERG/FLI1 inhibitor) was acquired from MedChemExpress and XRP44X (Ras-Net-Elk-3 inhibitor) from Sigma-Aldrich. The drugs were reconstituted in DMSO. MTS assay was used to determine viable cells in proliferation upon drug treatments with increasing concentrations at 72 h time point. REH cells (10000 cells/well) were seeded with drugs into 96-well plates with a final volume of 100 μl. Following drug treatment, cell proliferation was measured using CellTiter 96® AQ_ueous_ One Solution (Promega). 20 μl of CellTiter 96® AQ_ueous_ One Solution reagent per well was added and cells were incubated for 3 h in a humidified (atmosphere 95% air/ 5% CO_2_) incubator at 37°C. Absorbance was measured at 492 nm by a spectrophotometer (Thermo Scientific, Multiskan Ex). The background signal (no cells) was subtracted and the average signal from three technical replicate wells was used in calculations. In parallel, cell viability and count was measured based on Trypan Blue (Sigma-Aldrich) staining using Cellometer Mini Automated Cell Counter (Nexcelom Bioscience). Relative proliferation and cell amounts were calculated by normalizing to DMSO as a control sample.

### Visualization tools

Scatter plots in Figures 1–4 and heatmaps in Figures 1 and 3 were generated with Scanpy [110] and scVelo [31]. Regulon activity heatmaps in Figures 2 and 5 were generated with ComplexHeatmap [146]. Illustrations in Figures 1, 3 and 4 were created with BioRender [147]. Motif logos in Figure 2 were generated with HOMER [129]. Track plots from gene loci in Figures 4 and 5 were generated from UCSC Genome Browser [124]. Other plots in Figures 1–5 were generated using ggplot2 [148] and base R graphics [149].

## Declarations

### Ethics approval and consent to participate

This study was approved by the Regional Ethics Committee in Pirkanmaa, Tampere, Finland (#R13109) and conducted according to the guidelines of the Declaration of Helsinki. A written informed consent was received by the patient and/or guardians.

### Consent for publication

A written informed consent was received by the patient and/or guardians for publication of their data. Sensitive data will be stored in a controlled access database (EGA).

### Availability of data and materials

The datasets generated and analyzed in the current study are available in Gene Expression Omnibus under the accession number GSE148218 https://www.ncbi.nlm.nih.gov/geo/query/acc.cgi?acc=GSE148218 [130] and European Genome-phenome Archive under the accession number EGAS00001004374 https://www.ebi.ac.uk/ega/studies/EGAS00001004374 [131]. The Human Cell Atlas bone marrow scRNA-seq data was downloaded from https://data.humancellatlas.org/explore/proiects/cc95ff89-2e68-4a08-a234-480eca21ce79 [28]. Bulk ATAC-seq profiles of pro-B cells were acquired from GEO GSE122989 https://www.ncbi.nlm.nih.gov/geo/query/acc.cgi?acc=GSE122989 [52]. Data for GRO-seq in the REH cell line is available in GEO GSE67540 https://www.ncbi.nlm.nih.gov/geo/query/acc.cgi?acc=GSE67540 [55]. Additional data for ChlP-seq peak analysis was downloaded from GEO GSE45144 https://www.ncbi.nlm.nih.gov/geo/query/acc.cgi?acc=GSE45144 [132], GSE99019 https://www.ncbi.nlm.nih.gov/geo/query/acc.cgi?acc=GSE99019 [58] and GSE126300 https://www.ncbi.nlm.nih.gov/geo/query/acc.cgi?acc=GSE126300 [57]. Code related to analyses is available from GitHub [30] (https://github.com/systemsgenomics/ETV6-RUNX1 scRNAseq Manuscript 2020 Analysis).

### Author contributions

JM, ST, OL and MH designed the study. JM performed HCA and ALL BM scRNA-seq analysis. JM and ST performed scDD analysis. JM and MH designed the workflow for SCENIC and its benchmarking. AK participated in benchmarking scRNA-seq analysis. AK and ML performed ATAC-seq motif analysis. ML performed GRO-seq assays and processed the data. MB participated in NGS library preparation. MB, PIM and PR performed additional flow cytometry analyses to guide the data analysis. ST, SL and LO participated in collecting BM cells for scRNA-seq. RK and TL performed cell and library preparation for scRNA-seq, supervised by RL. ST, ML and AM performed the experiments with leukemia cell line models, supervised by OL, OS and MH. ST and SR performed flow cytometry and analyzed the results. ML performed drug sensitivity testing. MH, JM and ST wrote the manuscript and all authors commented on it.

## Acknowledgements

This study was supported by the Finnish Functional Genomics Centre (University of Turku, Åbo Akademi University), FIMM Technology Center Sequencing Laboratory (Biomedicum, Helsinki), Biocenter Finland and the Sequencing Service GeneCore Sequencing Facility (EMBL, Heidelberg, Germany) through providing sequencing services. The authors wish to acknowledge CSC - IT Center for Science, Finland and UEF Bioinformatics Center, University of Eastern Finland, Finland for computational, and Biocenter Kuopio FinGEEC for flow cytometry resources.

