## Supplementary figures for "Single cell characterization of B-lymphoid differentiation and leukemic cell states during chemotherapy in ETV6-RUNX1 positive pediatric leukemia identifies drug-targetable transcription factor activities"

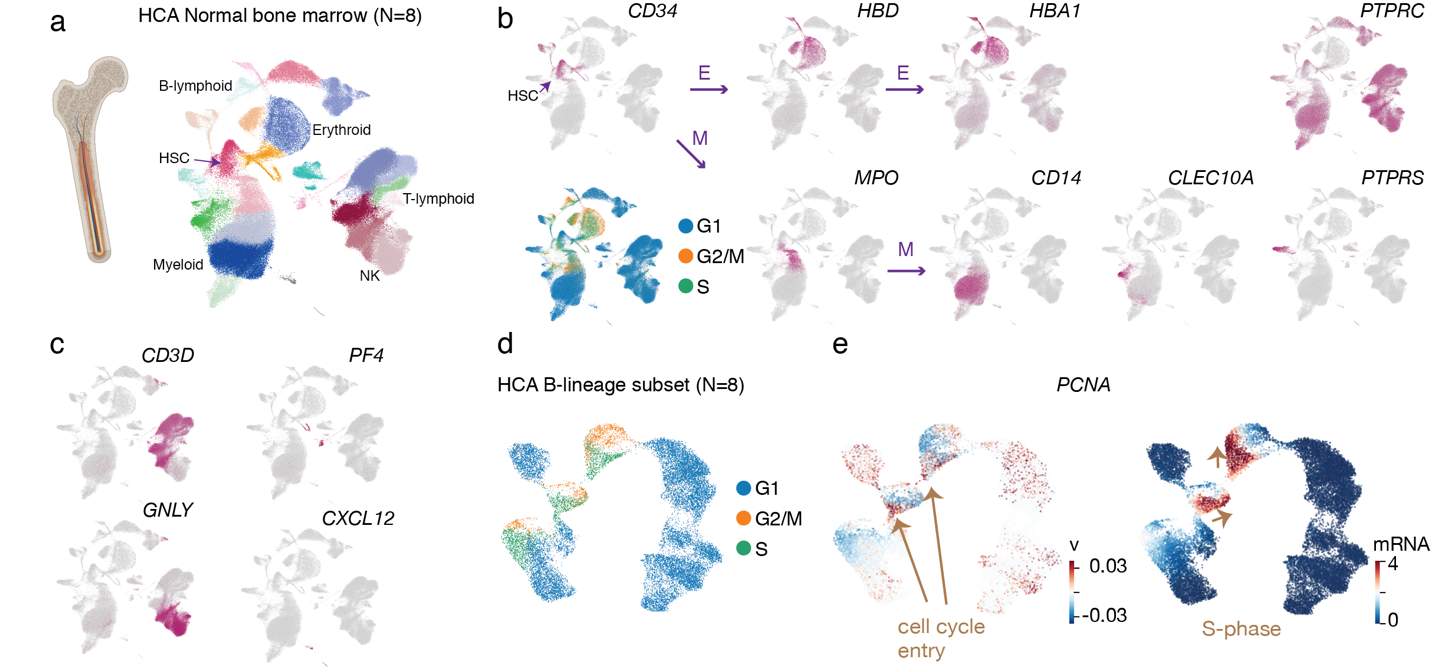

**Figure S1. HCA BM dataset**. a. Initial cell state assignment of Louvain clusters is presented on the two-dimensional UMAP visualization of HCA BM scRNA-seq (N=8). b-c. Expression level for classical lineage marker genes and the computationally predicted cell cycle state are visualized on the UMAP: stem cell (*CD34*), E: erythroid (*HBD*, *HBA1*), M: myeloid (MPO, CD14, CLEC10A for classical and PTPRS for plasmacytoid dendritic cells), T-lymphoid (*CD3D*), NK cell (*GNLY*), megakaryocyte (*PF4*) and *PTPRC* (also known as *CD45*) negative stromal (*CXCL12*) cells. d. Computationally predicted cell cycle state is shown on the HCA B-lineage subset UMAP. e. RNA velocity (left) and mRNA expression (right) for the S-phase marker *PCNA* are shown as in Fig. 1f.

Related to Fig 1.

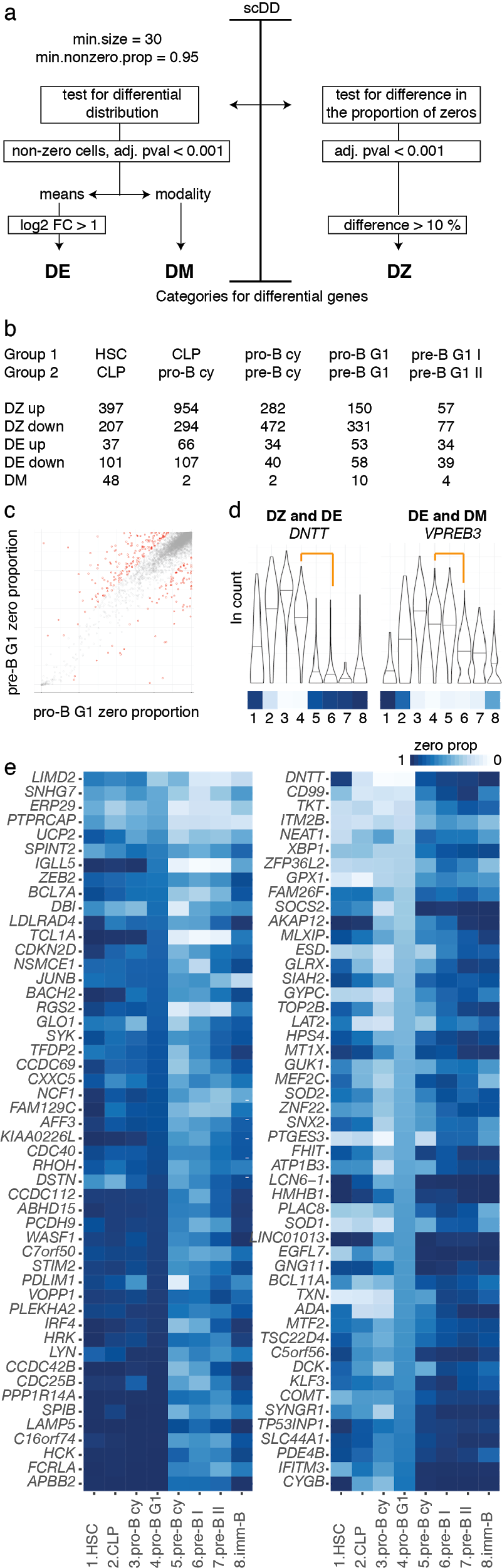

**Figure S2. Differential distribution testing between normal BM cell population using the scDD tool.** a. A schematic representation of filtering steps and cut-offs applied in the scDD analysis. See Methods for details. b. Number of differential genes along B-cell differentiation from HSC to preB-G1 II cell state is shown as a table. c. Scatter plot of ZP in pro-B G1 (x-axis) and pre-B G1 (y-axis) cell populations is shown. Red color indicates genes assigned to the DZ category. d. Violin plots for example genes showing the gene expression distribution found in the pro-B G1 to pre-B I comparison (refer to panel e numbering) in both DZ and DE analyses but not DM (*DNTT*), and in both DE and DM but not DZ (*VPREB3*). e. Top 50 genes up- and downregulated in pro-B to pre-B transition (in both G1 and cycling cell population comparisons) are visualized as a heatmap, where color corresponds to ZP. The darker color tones correspond to smaller portion of cells expressing the gene.

Related to Figs. 1, 3-5

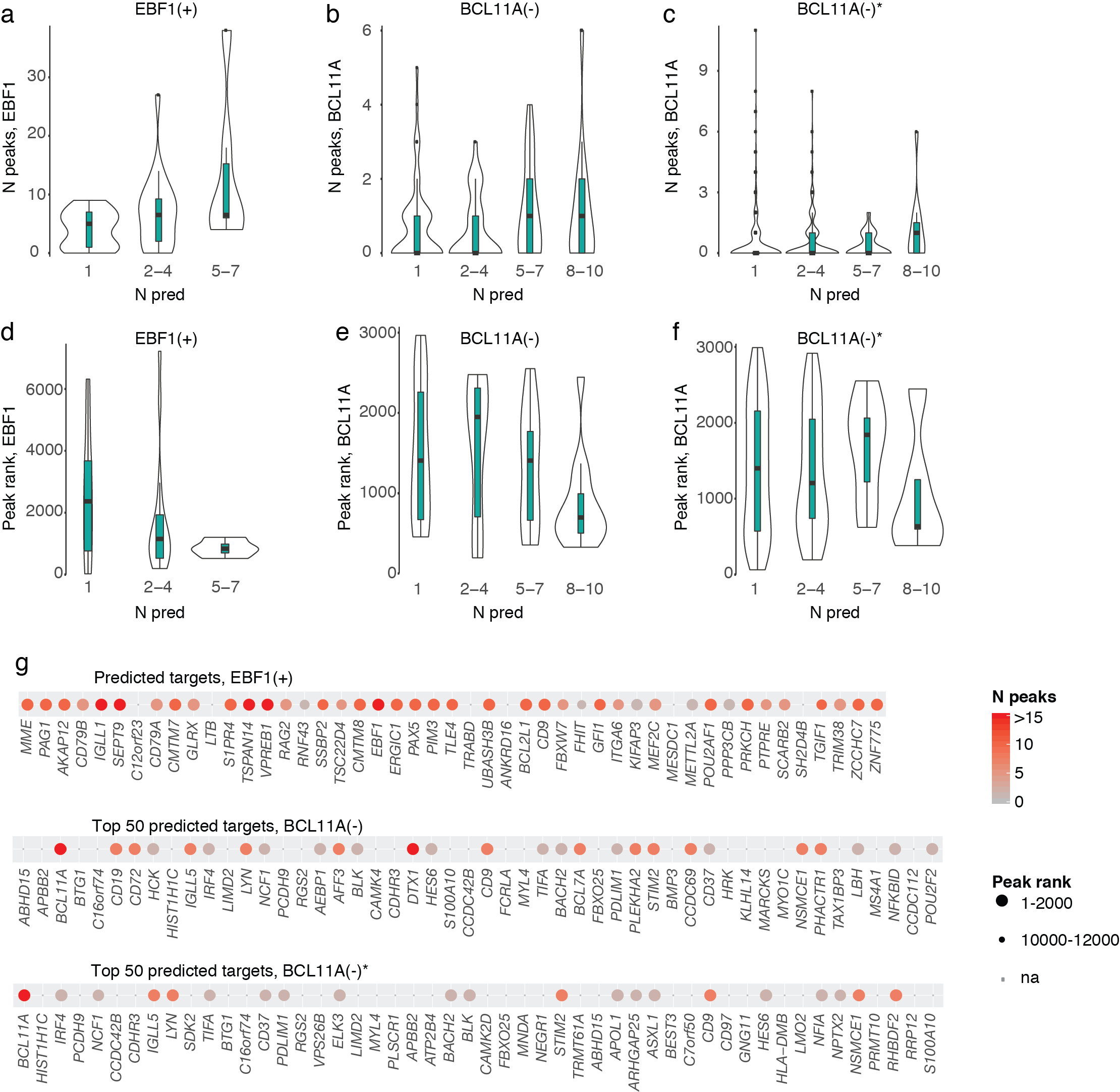

**Figure S3. ChIP-seq validation data for SCENIC regulons.** a-c. The distribution of ChIP-seq peaks associated to targets is shown as in Fig. 2e for EBF1(+), PAX5(+) and BCL11A(-) regulons obtained with the customized workflow. BCL11A(-)* corresponds to initial regulon discovered by default SCENIC run. d-f. The ChIP-seq peak rank distribution is visualized across binned regulon genes as in Fig. 2f. g. The ChIP peak data is visualized using a dot plot for top 50 predicted targets as in Fig. 2g.

Related to Fig. 2

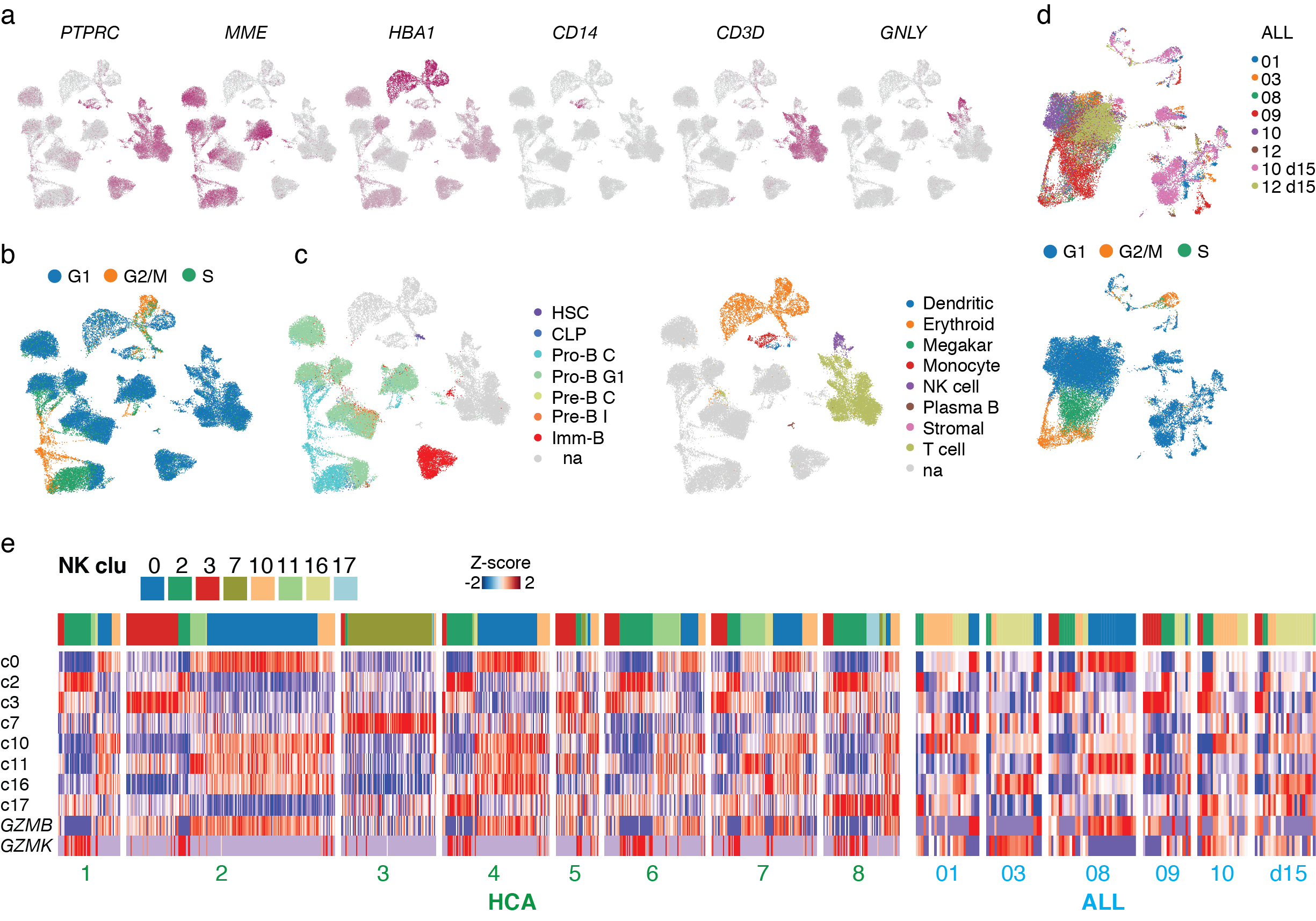

**Figure S4. E/R+ BM dataset.** a. Expression level for classical blood cell lineage marker genes is visualized on the UMAP: mature blood cell (*PTPRC*), B-lymphoid progenitor (*MME*) erythroid (*HBA1*), myeloid (CD14), T-lymphoid (*CD3D*) and NK cell (*GNLY*) cells. b. Computationally predicted cell cycle state is presented on the two-dimensional UMAP visualization of E/R BM scRNA-seq (N=8). c. Cell type assignment is shown for the E/R+ BM UMAP based on label transfer from HCA BM annotations (left: B-lineage cell states, right: other BM cell types). d. Sample origin (upper panel, donor in color) and computationally predicted cell cycle state (lower panel) are shown on the E/R+ BM batch-corrected UMAP (batch=donor). The largest cluster on the left (with cell cycle status G1 above and G2/M/S below) corresponds to leukemic cells.

Related to Fig. 3

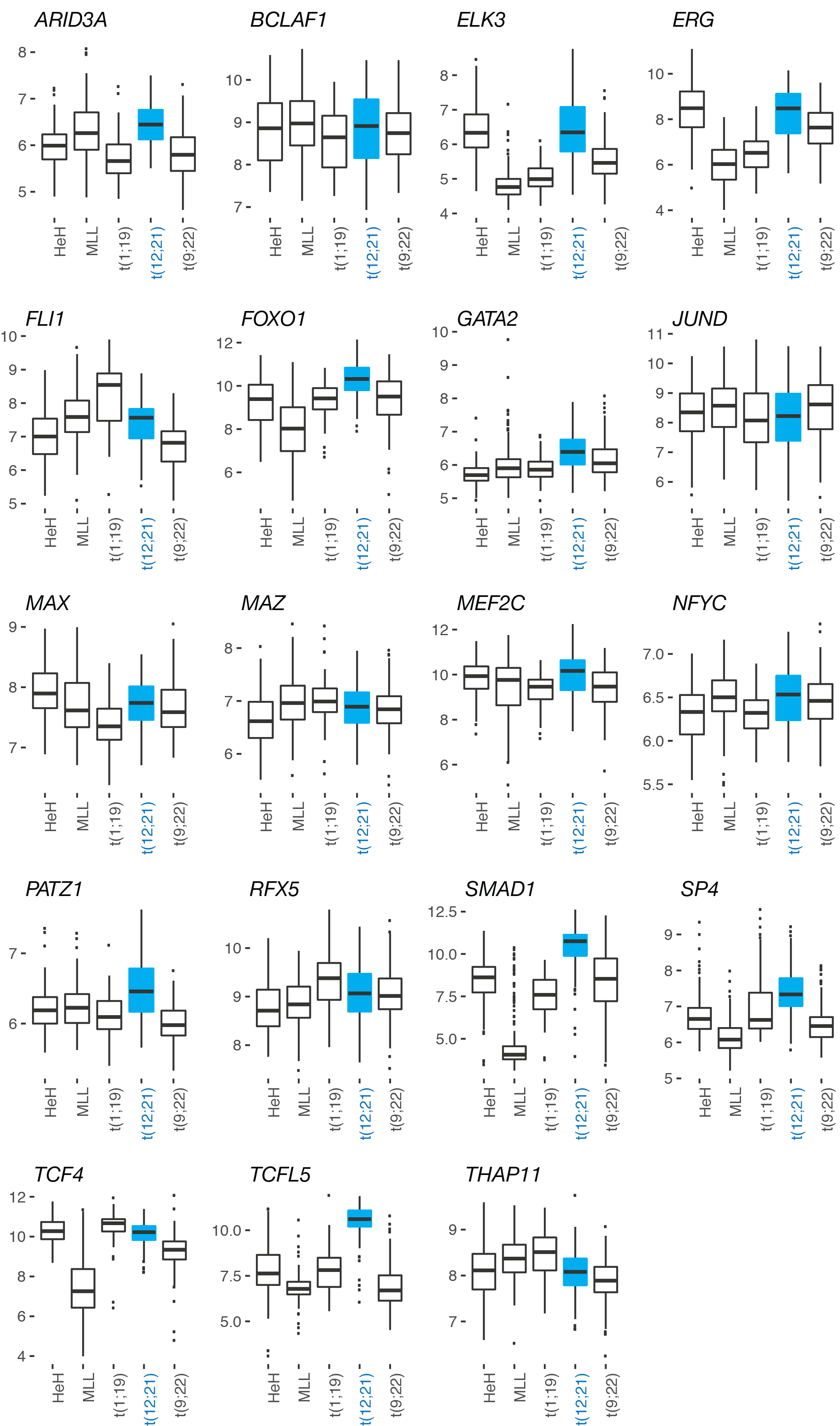

**Figure S5. Bulk gene expression data for pre-B-ALL subtypes**. The log2 gene expression levels in Hemap microarray dataset compared across pre-B-ALL subtypes as boxplots. TFs with high predicted TF activity in E/R+ ALL are shown.

Related to Fig. 5

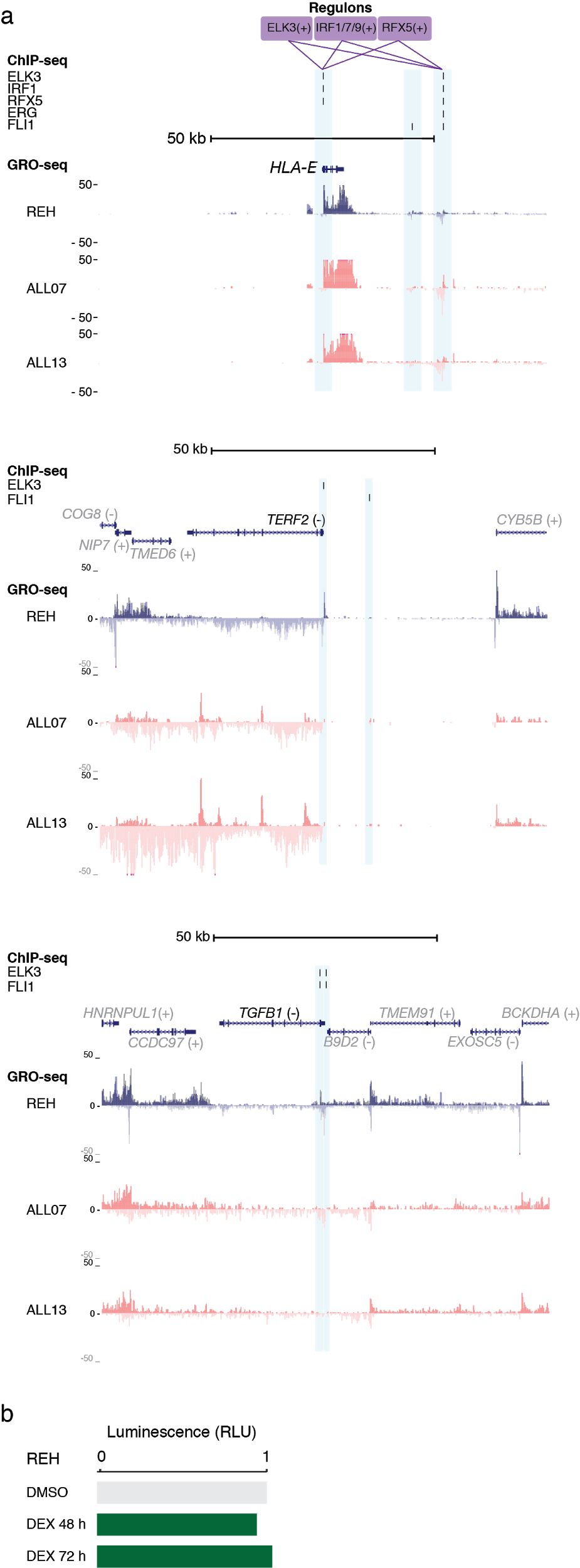

**Figure S6. Regulatory interactions associated with active TF regulons in E/R+ cells**. A. ChIP-seq and GRO-seq data are shown at the *HLA-E*, *TERF2* and *TGFB1* loci as in Fig. 5e. Active enhancers are highlighted by shading. The ChIP-seq peaks shown at these locations correspond to HUVEC (ELK3), HSC (ERG, FLI1), K562 (IRF1 upon IFNg stimulus) and GM12878 (RFX5) peak annotations. GRO-seq data is shown from E/R+ REH cell line and two primary E/R+ bone marrows. b. The luminescence signal from MTS assay in REH cells treated with the glucocorticoid dexamethasone (10 nM) is shown from 48 h and 72 h time points.

Related to Fig. 5
